# Does standard adjustment for genomic population structure capture direct genetic effects?

**DOI:** 10.1101/2024.05.03.592431

**Authors:** Ramina Sotoudeh, Sam Trejo, Arbel Harpak, Dalton Conley

## Abstract

Contemporary genomic studies of complex traits, such as genome-wide association studies (GWASs) and polygenic index (PGI) analyses, often use the principal components of the genotype matrix (PCs) to adjust for population stratification. In this paper, we explore the extent to which we may be discounting direct genetic effects by adjusting for PCs. Using family-based models that control for parental genotype (obtained via Mendelian imputation), we test whether PCs have a direct genetic effect on nine complex phenotypes in the White British subsample of the UK Biobank. Further, we assess the extent to which estimates of polygenic effects meaningfully change when adjusting for PCs in within-family models. Across the nine traits, within-family effects of the top 40 PCs are highly similar to their population effects, suggesting that standard PC adjustments diminish, albeit to a small degree, detectable signals of direct genetic effects. Within family models also confirm that PCs have significant marginal effects on a few traits, most consistently, height and educational attainment. Nonetheless, the variance explained by the effects of PCs is modest, and adjusting for PCs does not appear to affect the magnitude and significance of PGI effects in within-family models.

## Introduction

Since at least 2006 (see Price et al. 2006), including principal components (PCs) of the genotype matrix as covariates has become a popular method for controlling for population stratification when conducting genome-wide association studies (GWAS) and polygenic index analyses—replacing the genomic control approach (Devlin & Roeder 1999). More recent methods advocate the use of mixed models to account for pairwise genetic relatedness in the sample (Price et al. 2010; Zhou and Stephens 2012; Vilhjálmsson & Nordborg 2013). New methods for conducting GWAS in multi-ancestry samples also utilize PCs to detect ancestry-specific effects (Wojcik et al. 2019).

Principal components have a number of desirable features, including their ease of computation and their tendency to correspond to the geographic distribution of populations (Novembre et al. 2009; Urnikyte et al. 2019; Chen et al. 2009; Wang et al. 2012). As such, PCs have gained popularity in the population genetics literature. But the extent to which adjusting for PCs also attenuates direct genetic effects is unknown. Here we ask, how much of the direct genetic signal is lost when we adjust for PCs across phenotypes? That is, there may be alleles that display systematic differences across major axes of ancestry (and thus evince high eigenvalues on PC vectors) with direct genetic effects.^1^ While the top genetic PCs tend to correlate strongly with the distribution of individuals across environments, we might expect some causal signal to also be captured by these axes of variation among subgroups of people (whether because of genetic drift, assortative mating, or selective pressures). Changes in environmental effects, in fact, may induce genetic selection to restore optimal fitness; this would, in turn, lead to changes in allele frequency along the same dimensions as the ‘confounding’ environmental difference—i.e., the PC space (Harpak and Przeworski 2021; Mills and Mathieson 2022). In other words, direct genetic effects—to the extent they emerged in response to the environment—may lie along the very PCs that we purge as confounds in the typical GWAS.

With this in mind, the present paper evaluates whether there is signal in genetic PCs that has direct effects on certain phenotypes. We examine both additive and (additive-by-additive) interaction effects. We then assess whether our estimates of polygenic effects in within family models meaningfully change when controlling for the direct effects captured by PCs. We consider nine complex phenotypes in our analyses, including: body mass index, birthweight, number of cigarettes smoked per day, diastolic blood pressure, drinks per week, height, number of children ever born, having seen a general practitioner (GP) or a psychologist for depression and number of years of schooling completed (following the coding scheme in Okbay et al. 2022) which we henceforth refer to as educational attainment.

## Background

Genetic principal components are commonly used as indicators of genetic population structure (McVean et al. 2009; Patterson, Price and Reich 2006). Principal components, in general, are vectors that result from a dimension reduction technique meant to reduce the dimensionality of the data while retaining most of the information (Hotelling 1933).

When applied to molecular genetic data, principal component analysis maximizes the allelic variance that can be captured in a given number of variables (or components), under the constraints that the variables are: 1) linear combinations of the underlying allele counts and 2) linearly independent of each other. For example, in the White British subsample of the UK Biobank, the first PC tends to distinguish between those born in England versus those born in Scotland and the second PC tends to distinguish between those born in Wales versus those born in England or Scotland (see Figure S1, SI). Higher order PCs capture less interpretable, and often less replicable axes of ancestry. Young et al. (2019), for example, show that of the 40 PCs assessed, PCs beyond the top seven in the White British UKB sample do not replicate in a held-out sample, suggesting that they are capturing noise and local, same-chromosome linkage disequilibrium (LD) rather than population structure.

PCs have also become widely used in population genetics research to adjust for population stratification (see e.g., Mathieson et al. 2023; Akimova et al. 2023; Okbay et al. 2016; Okbay et al. 2022; Mills et al. 2021; Duncan et al. 2019). Population stratification refers to when population structure (i.e., differences in allele frequencies within different subpopulations) causes bias in genetic analyses due to its correlation with phenotypic differences through genetic background and/or environmental confounding (Vilhjálmsson & Nordborg 2013). Not properly accounting for population stratification can lead to results that overstate the role of genetic effects, which become confounded by environmental factors (or even other genes through long-range LD) (Price et al. 2006; Hamer 2000).

By design, the top PCs will tend to load on SNPs that reveal the genetic structure of a population—that is, SNPs that occur at relatively higher or lower frequencies in different subpopulations. This will be true regardless of whether the SNP in question has a direct effect on an outcome of interest. Adjusting for PCs may therefore unintentionally discount the effects of causal SNPs when those SNPs vary in frequency between subpopulations. In this paper, we ask two key questions: 1) whether there is direct genetic signal captured by PCs and 2) if including PCs in our models affects, in any meaningful way, conventional polygenic prediction estimates (i.e., from polygenic indices calculated based on population GWASs).

## Analytic Strategy

Simply regressing various phenotypes on the top genetic PCs in a sample of unrelated individuals is problematic. By picking up population structure, PCs also capture environments shared by those in a given subpopulation—everything from differences in inherited wealth by region (Townsend 1979) to biased treatment by teachers (Bishop et al. 2005) or employers (Levon et al. 2021) based on characteristics associated with different regions of the UK, such as, say, an accent. Indeed, this is precisely the reason that PCs are used as covariates in GWAS and PGI analyses: to regress out the confounding influences of environments that are correlated with genotypes, as well as of correlated markers across the genome due to linkage disequilibrium (Price et al. 2006). Because PCs pick up population structure, however, straightforward estimation of their effects on phenotypes is difficult.

Our approach therefore rests on first purging, as much as possible, the possibility of confounding from our models before estimating the direct effects of PCs. We do so in two ways. First, we deploy PCs computed in a relatively homogenous sample by limiting our analyses to the White British subsample of the UK Biobank (UKB). Since siblings with parents of more distinct ancestries are likely to have higher discordance on their PCs (as well as higher heterozygosity rates), the results may be driven by these individuals in our sample, who may differ from others on unobserved characteristics. We remove them to ensure that no such dynamics are driving the results.

However, even with a relatively homogenous sample, population stratification may still be at play; for example, people of Scottish and English ancestry in the UK have both different average physical and cultural environments and different allele frequencies. We therefore run within-family models to minimize the extant association of confounding factors. Family-based models are generally accepted to yield genetic estimates that are less susceptible to environmental and genetic confounding (Veller & Coop 2023). Siblings share the same parental gene pool (as well as cultural patrimony and family environment), meaning that genetic differences between them—whether at a single locus or summarized via a PGI or PC—is the result of the random assignment of alleles during meiotic recombination and segregation. Thus, estimating the effects of PCs using within-family models provides estimates of direct genetic effects that are not affected by environmental confounds.^2^ However, while less susceptible to bias than population-based analyses, family models can still be susceptible to genetic confounding on a given chromosome – i.e. linkage of non-local but same-chromosome loci due to assortative mating or selection (Veller & Coop 2023) and the extent to which their estimates can be generalized to the population of interest is unclear (Fletcher et al. 2024; Veller, Przeworski & Coop 2023)^3^.

Traditional approaches to within-family analysis rely on sibling models which, in effect, regress sibling differences in phenotype on sibling differences in polygenic index. While effective at removing environmental confounding, these approaches can suffer from low statistical power.^4^ Leveraging Mendelian randomization within family in order to impute missing genotypes, by contrast, has been shown to increase power by increasing the effective sample size (Young et al. 2022). Specifically, we use Young et al.’s method (2022) for non-linearly imputing missing parental genomes using genotyped sibling pairs. Because each non-twin sibling’s autosomal genotype in a family is effectively a random draw from the parents’ genotypes, it is possible to use observed sibling genotypes to work backwards and partially recover parental genotypes.

In the UKB, families vary in the missingness of parental genotypes. In cases where both mother and father are missing, Young et al.’s approach infers the same genotype for each parent. To ensure that the results are applicable to all patterns of parental missingness, we calculate the mid-parent genotype – i.e., the average genotype of the two parents – for each family. When both parents are missing, the mid-parent genotype is the same as the average of each of their individual imputed genotypes. When one parent is missing, mid-parent genotype is the average between the imputed parent’s genotype and the observed parent’s genotype. And when both are present in the data, it is the average of their observed genotypes.

We calculate PGIs for nine different phenotypes based on the most recent population-based GWASs for each phenotype and the first 40 genetic PCs for both siblings and mid-parents. This, in turn, allows us to condition on the mid-parental PGI and PCs in our various regression models. The effect of a child’s PGI or PC can consequently be interpreted as a direct genetic effect, above and beyond any environmental information captured by the parental controls.

## Data and Measures

Our data come from the UK Biobank, a national biomedical database collected in the United Kingdom that links participants genotypes to a broad questionnaire about their physical and mental health, behaviors, and demographics. The full UKB has more than 500,000 respondents from a wide range of social and ethnic backgrounds, including a subsample of 408,219 respondents who are White British (see Bycroft et al. 2018 for how these individuals were identified). For most of our analyses, we focus on the subset of respondents who have at least one parent or sibling in the data and were identified as White British (n = 38,738) following the procedure described below.

To prepare the genetic data, we applied a standard protocol for genetic quality control. We restricted the genotype data to SNPs with a minor allele frequency of at least 0.05, filtered out individual genotypes with high rates of missingness (>10%), removed SNPs that violated the expectations of the Harvey-Weinberg equilibrium (p-value of HWE test < 1e-6), and dropped individuals who had a mismatch between their self-reported sex and their XY chromosome composition.

To identify related individuals, we utilized the KING software package. KING infers identity-by-descent information using SNP data, thereby allowing unrelated individuals to be reliably separated from related individuals of varying degrees. Monozygotic twins, half siblings, and first, second, and third-degree cousins were removed from the analyses. We then distinguished sibling pairs and parent-child pairs (who we henceforth refer to as our sibling and family sample) from the unrelated individuals remaining in the data. Accurately estimating genetic PCs depends on having first removed cryptically related individuals from the genotype matrix. For this reason, although our core analyses focus on the sibling and family sample, PCs were first estimated on the sample of unrelated White British individuals. We then project individuals from the sibling and family sample into this PC space.

More specifically, we calculated the genetic PCs for sibling and mid-parent genotypes in three steps. First, we used the bigsnpr package in R to clump SNPs that are in linkage disequilibrium (LD), remove long-range LD regions, and then estimate the first 40 PCs for all White British unrelated individuals in the UKB (Bycroft 2018; Privé et al. 2018). We then identified outliers along these 40 PCs, removed them from the data, and recalculated the PCs using this more homogenous set of unrelated individuals. Finally, we projected the sibling and mid-parent genotypes onto this PC space to obtain their loadings along these 40 PCs. By projecting our siblings and family sample onto the PC space of the unrelated individuals, we ensure the PCs pick up broader genetic structure in the White British population (rather than artifactual genetic structure mechanically present in close relatives).

We calculated PGIs in two different ways. The first approach mimics how PGIs are traditionally constructed in the literature – using population GWASs. We first identified summary statistics from the largest available GWAS of each phenotype: height^5^ (Yengo et al. 2022), birth weight (Horikoshi et al 2016), number of cigarettes per day (Liu et al. 2019), diastolic blood pressure (Surendran et al 2020), drinks per week (Liu et al 2019), years of educational attainment (Okbay et al. 2022), number of children ever born (Mathieson et al. 2023), and depression (Nagel et al. 2018).

As part of standard quality control measures, these GWASs exclude cryptically related individuals, including, importantly, siblings. Thus, while many of these GWASs include the UKB, they excluded the sibling subsample that we use here. We then used PRScs to adjust the GWAS summary statistics to account for the linkage disequilibrium structure of the UKB sample. Finally, we used the PGI construction script provided by Young et al. (2022) to construct PGIs for probands (siblings) and the mid-parent in each family.

While using the common approach to PGI construction makes our results directly applicable to previous work, a potential issue with this approach is that population-based GWASs can vary in the homogeneity of their samples and how many PCs are adjusted for. We therefore also conducted our own GWAS for each phenotype using a sample of unrelated White British individuals in the UKB as our discovery sample, adjusting for the first 40 White British PCs. By comparing these two different approaches, we can be sure that any differences we see in the results are not due to different choices in how population stratification is accounted for at the GWAS stage. The steps for constructing the PGI after having run our GWASs are the same as in the traditional approach that uses external GWASs. The results from using our own GWAS can be found in the SI in Table S4.

## Model Specifications

To evaluate the direct genetic effects of PCs and how their inclusion in models affects PGI estimates, we ran three different sets of regression models.

We refer to the first set of models as within-family PC models. In these models, we directly evaluate the effect of PCs on our nine phenotypes while adjusting for mid-parent PCs to account for family structure. We begin by defining a base model that includes the first 40 mid-parent PCs, as well as the age, sex and genotyping array of the proband. We then incrementally add proband PCs in groups of four to the model and evaluate how their addition affects variance explained by the model (R^2^), using an F-test to determine whether any incremental gains in R^2^ are statistically significant.

The base model can be formalized as the following, for individual *i* in family *j*:

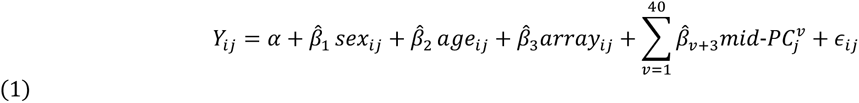

Where *Y*_*ij*_ is the phenotype, 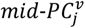 is the *v*^*th*^ midparental genetic PC, α is the intercept term, 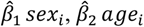, and 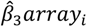 are the effects of our proband controls for sex, age, and array respectively, and ϵ_*ij*_ is the error term.

To this base model, we add the proband genetic PCs (where 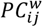 is the *w*^*th*^ genetic PC) in groups of four. For example, in the first set of models building on the base model, we add PCs 1-4:

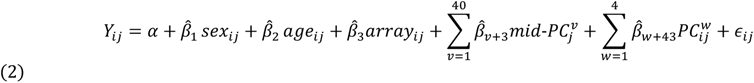

The model with probands 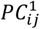 through 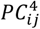 shown in equation 2 is then compared to the base model in equation 1 using incremental R^2^ and an F-test. In subsequent regressions, we continue to add the next four proband PCs until all 40 proband PCs are in the model. For example, the third model includes proband 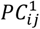 through 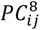:

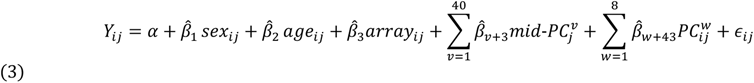

As we add additional proband PCs, we always compare the new model to the previous model and not the base model. Thus, in the examples above, the third set of regressions (equation 3) are compared with the second (equation 2).^6^ This ensures that our incremental R^2^ results always capture the gains to explained variance that result from adding more proband PCs.

### Interaction effects

In addition to the additive models of PCs reported above, we also examined whether PCs exhibit interaction effects for the different phenotypes considered here. While GWAS and PGI analyses typically adjust for main effects of PCs, they do not include non-linear functional forms. So, this portion of the analysis does not inform the question of whether including PCs leads to attenuation of direct genetic effects in GWAS or PGI models but rather asks if dimensions of population structure interact in ways that impact phenotypes through direct genetic effects.

The number of potential cross-PC combinations is extremely large (considering just the possible bivariate combinations between the 40 proband PCs would lead to 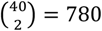 regressions). We therefore focus exclusively on bivariate interactions between the first eight PCs, in addition to squared terms for each PC to test quadratic effects for our nine phenotypes. We also include the bivariate interactions between each PC and the phenotype’s relevant PGI.

Here, for each bivariate combination, we compare the amount of additional variance that is explained when moving from an additive to an interaction model. That is, we compare a purely additive version of each model, where the relevant proband PCs, parental PCs and proband controls were included, to a model where the proband PCs are also interacted. For example, in the case of 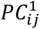 and 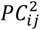, the model fit of equation 5 was compared to that of equation 4:

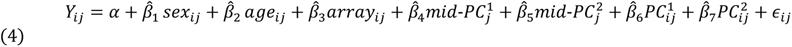

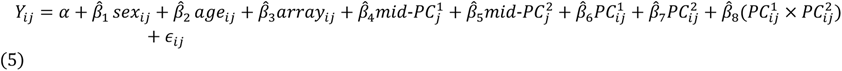

Unlike the additive models, these models are not cumulative (i.e. lower order PCs are not included in the model when higher order PCs are examined). In the case of quadratic effects, instead of two PCs, we adjust for a given PC (for the mid-parent and the proband), and the interaction term is a squared term of that particular PC for the proband.

### Changes in PGI estimates

Finally, we assess whether the effects of PGIs (based on population GWASs) on a phenotype meaningfully change when we add proband PCs to within-family models. We use a test from Clogg, Petkova, and Haritou (1995) for evaluating the significance of changes in coefficient magnitudes between nested models. If we are concerned about the attenuation of direct genetic signal, we should see the magnitude of PGI estimates significantly change once proband PCs are included in within-family models.

Within-family models are less prone to confounding (with the caveats discussed above). Still, PGIs constructed with weights from population GWASs contain both environmental and genetic confounding, and even within-family models cannot be interpreted as only including direct genetic effects. Including PCs in the model may affect the PGI models either through absorbing direct genetic effects, or residual confounding that exists in the PGIs. Here we ask whether the PGI estimates meaningfully change when accounting for the direct effects of PCs on the phenotypes by absorbing direct, indirect, or confounding effects present in the PGI. We chose to use population-level PGIs to make these analyses relevant to the largest number of researchers possible. Although sibling GWASs continue to increase in size, they still lack power, and thus many researchers opt for better-powered GWASs conducted on population samples.

Again, we compare a model with proband PCs to a base model without them. The base model is as follows:

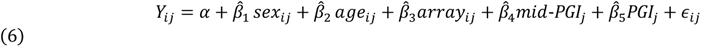

Where *Y*_*ij*_ is the phenotype, *PGI*_*ij*_ is the proband’s PGI for the relevant phenotype, *mid-PGI*_*j*_ is the mid-parent PGI for the phenotype. Sex, age, and the genotyping array of the proband are included as controls.

To this base model, we add the PCs of the proband in groups of 4. The models are again cumulative, comparing each model to the version before it. For example, the second set of models, shown in equation 7 is compared to the base model in equation 6. To assess the total impact of adding proband PCs to the model, we also compare the PGI estimate from the final model with all 40 proband PCs to that of the base model.

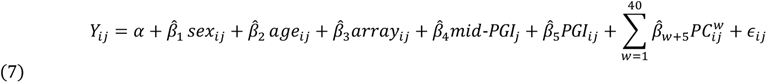

Because we are running dozens of regressions per phenotype, we use the Benjamini-Hochberg approach (BH) to adjust p-values in order to account for the likelihood of Type I errors (false discovery rate of 5%). Within a set of models (additive, interaction, PGI), we rank the p-values for the main coefficients of interests from the different regressions in ascending order. We then multiply each p-value by the total number of regressions and divide by its rank (Benjamini and Hochberg, 1995).

## Results

We first report results from the additive models examining the direct effects of proband PCs on our phenotypes of interest. In Figure 1, we report incremental R^2^ values across different model specifications. This provides a sense of how much of the overall variance in these phenotypes is explained by the first 40 proband PCs, parental PCs, and proband and parental PCs combined. The amount of variance captured by PCs (proband, parental and both) varies meaningfully by phenotype. We use an ANOVA to test whether adding PCs significantly improved variance explained. The results show that full models (age + sex + array + 40 parental PCs + 40 proband PCs) explained significantly more variance than a model including only age, sex, array, and 40 parental PCs for height, educational attainment, BMI and number of children ever born (as indicated by asterisks in Figure 1). The magnitude of these improvements, however, are small, suggesting that PCs capture only a small portion of the variation in these phenotypes (Table S1 in the SI reports the R^2^ values that are portrayed in Figure 1).

**Figure 1:**
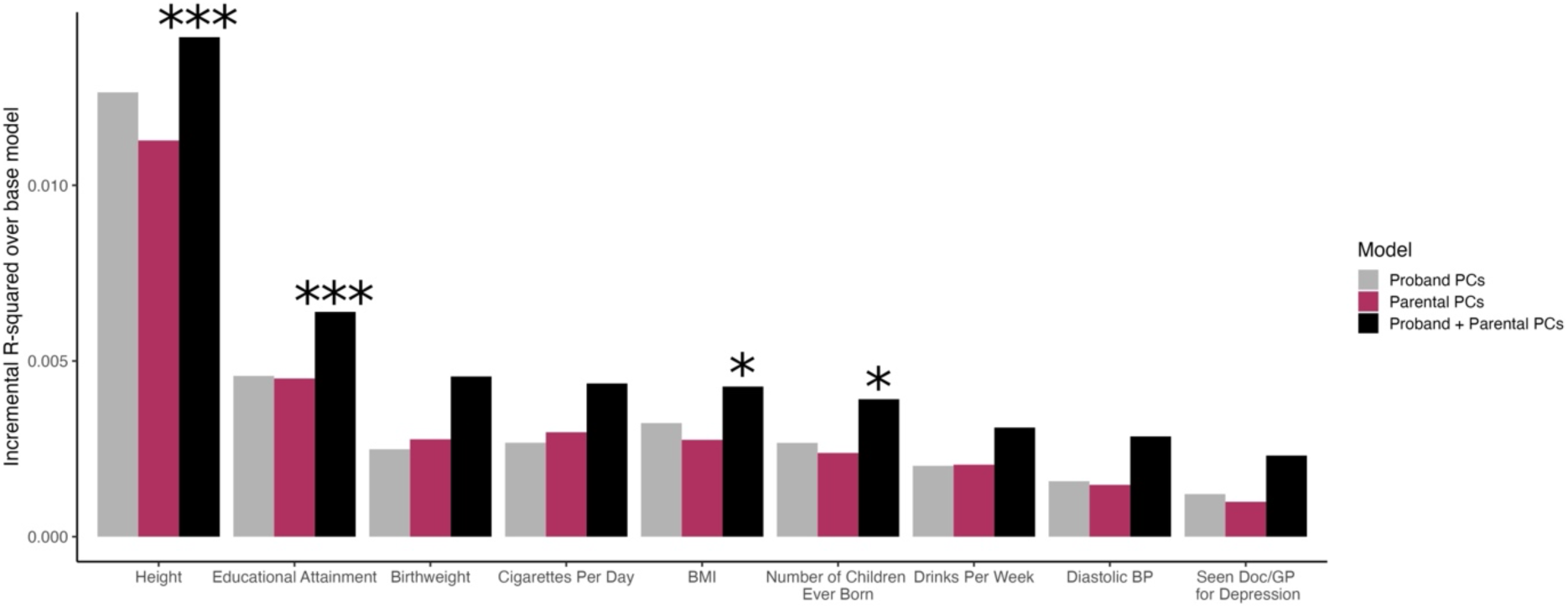
Incremental R^2^ across different model types and phenotypes. Incremental R^2^ over the base model containing only age, sex, and array is reported across different model specifications. An ANOVA was used to compare the full model (age + sex + array + parental PCs + proband PCs) to the model with age + sex + array + parental PCs. Asterisks reflect the degree of statistical significance for the ANOVA: * p < 0.05, ** p < 0.01, and *** p < 0.00. R^2^ values used to produce this figure are reported in Table S1 in the SI.

We also conducted a set of ANOVAs to compare models with only age + sex + array to ones where we add 40 proband PCs and 40 parental PCs separately. Compared to the base model (age + sex + array) adding 40 parental PCs returned a significant ANOVA test for all phenotypes except depression and diastolic blood pressure. Similarly, adding 40 proband PCs was significant for all phenotypes except depression. For the sake of visual clarity, these tests are not shown in Figure 1.

Figure 2 plots within-family effects of PCs by effects in population models. Within-family models include White British siblings, and adjust for 40 parental PCs and age, sex, and array as controls. Population models include unrelated White British individuals, and do not adjust for parental PCs. Overall, the estimates of within-family and population models are largely similar, suggesting that PCs in population models are, at least in part, capturing direct genetic effects.

**Figure 2:**
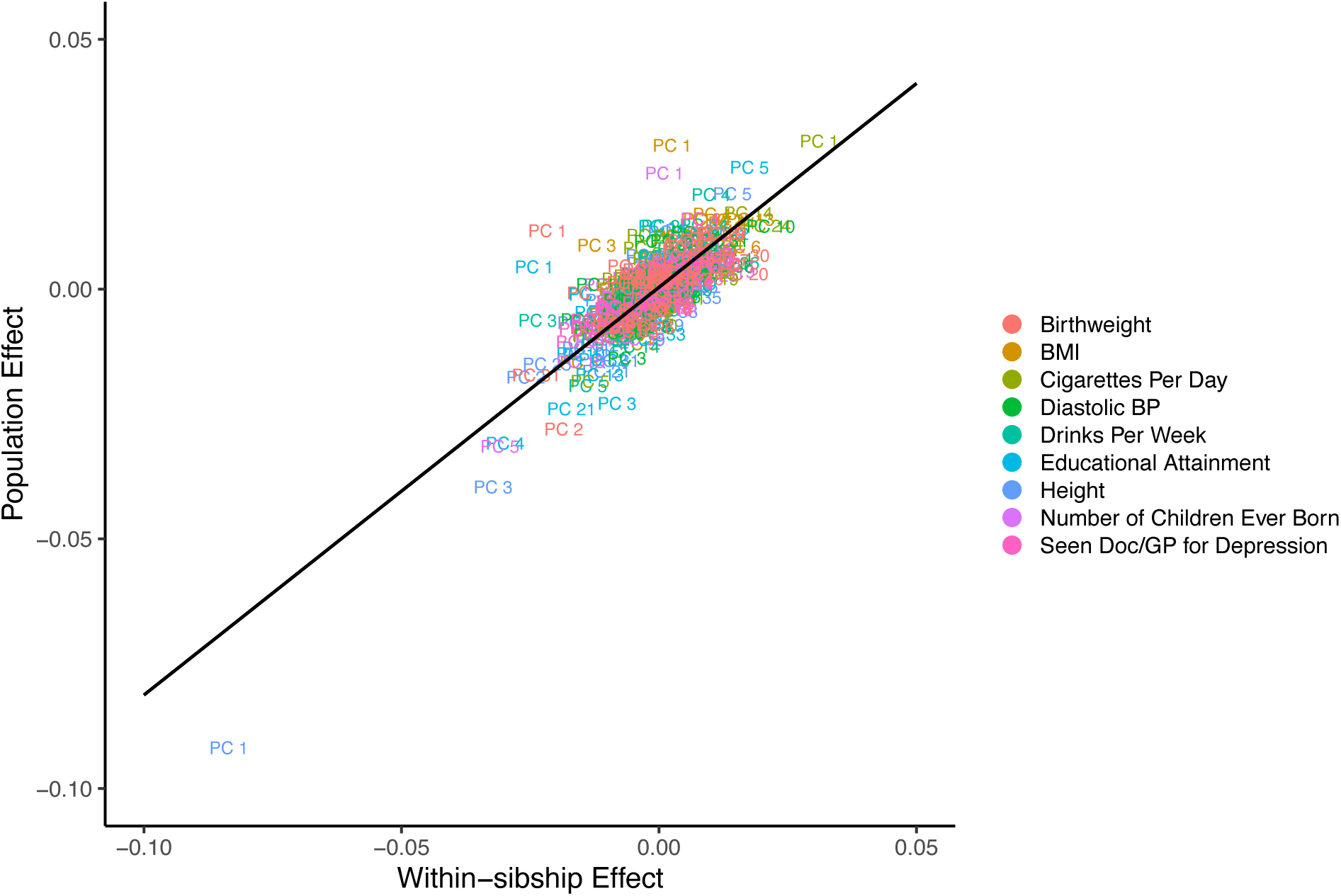
Comparing within-sibship effects and population effects of PCs, across phenotypes. Within-sibship versus population effects of 40 PCs on phenotypes (in standard deviation units) are plotted along with the best fit regression line. The population models include controls and all 40 proband PCs and use the unrelated White British subsample. The within-family models use the White British sibling subsample and include controls, all 40 proband PCs, and all 40 parental PCs. Across phenotypes, effects of PCs for the two different models are highly overlapping, with the exception of some lower order PCs (1-5).

Figure 3 plots the point estimates from each model ∓ 95% confidence interval. Within-family effects are shown in orange and population effects are shown in black. The confidence intervals are not BH corrected, and results are meant to help us descriptively assess the extent to which the direct effects and population effects of PCs overlap. We include the first 8 PCs for brevity; Figure S2 in the SI includes all 40 PCs.

**Figure 3:**
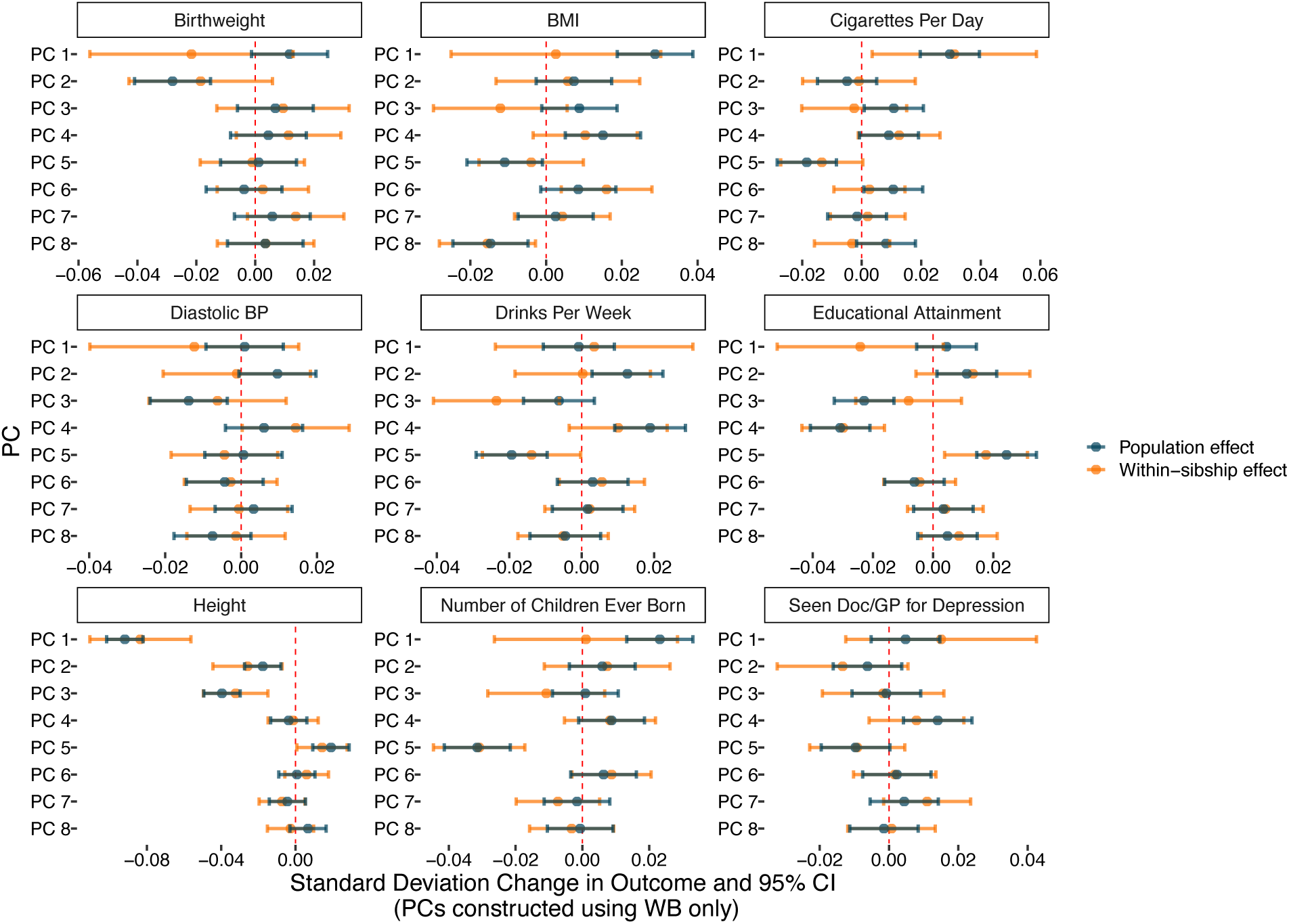
Comparing within-sibship effects of PCs to their population effects, by phenotype. Standard deviation change in phenotype by PCs (point estimate ∓ 95% confidence interval) are plotted in population models (black) and in within-family models (orange). The nine panels correspond to the nine different phenotypes included in this study. The population models include controls and all 40 proband PCs and use the unrelated White British subsample. The within-family models use the White British sibling subsample and include controls, all 40 proband PCs, and all 40 parental PCs. For the sake of visual clarity, the first 8 PCs are plotted here; Figure S2 in the SI plots all 40 PCs.

Here too, we see that within-family and population estimates are highly overlapping, though the magnitude of the effects are small. For example, PCs 1, 2, and 3 predict height in both within-family and population models. In the few exceptions, such as PC 1 for number of children ever born and BMI, PCs are only predictive in population models, and not in within-family models, suggesting that PCs for these phenotypes are capturing population stratification in population models.

Next, we turn to examining the incremental gain in R^2^ that results from adding different numbers of proband PCs to the models. Table 1 reports the incremental R^2^ of the models and the p-values that obtain after applying BH correction. Recall, the relevant comparison model for each model is one that has four fewer proband PCs in it. The results are reported in ascending order according to their BH-corrected p-values and we only report models that reach statistical significance after BH correction.

**Table 1:** Significant effects of PCs in additive models.

| Phenotype | PC set | Incremental R<sup>2</sup> (%) | BH corrected p-value |
| --- | --- | --- | --- |
| Height | PCs 1-4 | 3.053 | 0 |
| Educational Attainment | PCs 1-4 | 2.351 | 0.005 |
| Number of Children Ever Born | PCs 5-8 | 1.798 | 0.006 |
| Birthweight | PCs 29-32 | 3.077 | 0.036 |

**Table 2:** Significant interaction effects of PCs.

| Phenotype | PC set | Incremental R<sup>2</sup> (%) | BH corrected P-value |
| --- | --- | --- | --- |
| Educational Attainment | PC 2 x PC 2 | 3.75 | 0.00002 |
| Height | PC 3 x PC 4 | 1.90 | 0.00003 |
| Height | PC 2 x PC 2 | 1.73 | 0.0001 |
| Educational Attainment | PC 1 x PC 2 | 2.60 | 0.0005 |
| Height | PC 2 x PC 4 | 1.37 | 0.0009 |
| Drinks Per Week | PC 1 x PC 2 | 1.16 | 0.0012 |
| Educational Attainment | PC 1 x PC 5 | 2.05 | 0.0021 |
| Seen Doc/GP for Depression | PC 2 x PC 4 | 1.53 | 0.0029 |
| Birthweight | PC 4 x PC 4 | 3.20 | 0.0040 |
| Educational Attainment | PC 5 x PGI | 0.65 | 0.0059 |
| Number of Children Ever Born | PC 1 x PC 3 | 1.06 | 0.0149 |
| Educational Attainment | PC 3 x PC 3 | 1.43 | 0.0183 |
| Diastolic BP | PC 2 x PC 2 | 0.96 | 0.0183 |
| Drinks Per Week | PC 2 x PC 2 | 0.73 | 0.0186 |
| Diastolic BP | PC 1 x PC 5 | 0.91 | 0.0216 |

After BH correction, only four phenotypes exhibit significant effects for PCs. Height and educational attainment see significant improvement of model fit when the first four PCs are included, number of children ever born sees significant improvement in model fit once PCs 5-8 are added to the model, and birthweight sees significant improvement when PCs 29-32 are added to the model.

Based on these results, PCs with significant effects on phenotypes appear to be predominantly lower order (PCs 1 through 8). Figure S2 in the SI provides additional evidence – across all nine phenotypes, 13 out of the 37 significant hits in within-family models (before BH correction) come from the first 8 PCs and 26 come from the first 20 PCs. By implication, if attenuation of direct genetic effects by including PCs is indeed a concern, including a smaller number of PCs should do very little to counter it.

### Interaction Effects of PCs

We next examine interaction effects. Incremental R^2^ values and BH-correct p-values are reported in Table 3. As mentioned above, interaction models are compared against an additive base model with the same parental genetic variables as well as the proband’s added rather than multiplied.

Quadratic effects are small but present for educational attainment, height, diastolic blood pressure, drinks per week and birthweight. Interestingly, most of these quadratic effects are due to PC2^2^, the exceptions being birthweight (PC4^2^) and a second finding for educational attainment (PC3^2^). We also find evidence for interaction effects between PCs, most notably among PCs 1 to 5 for height and educational attainment. For educational attainment, we find significant effects after BH correction when PC 1 and 2 are interacted, when PC 1 and 5 are interacted, and when PC 5 is interacted with the PGI. For height, we see significant effects when PC 3 and 4 are interacted and when PC 2 and 4 are interacted. We also find significant associations with drinks per week (PC1 x PC2), depression (PC2 x PC4), number of children ever born (PC1 x PC3), and diastolic blood pressure (PC1 x PC5).

### PGI models

Finally, we examine whether PGI (based on the largest available population GWAS) estimates are affected when proband PCs are included to our additive models. The results are reported in Table S3 of the SI. Across the board, we find that the addition of PCs does not substantially or significantly change coefficient estimates. These results imply that while PCs may capture some modest direct genetic effects, we are not attenuating our PGI estimates by adjusting for them. We find similar results using PGIs from standardized GWASs that we constructed ourselves using the UKB unrelated population (more information on those GWASs can be found in the SI).

There, again, we find no significant or substantial differences in PGI estimates before and after the inclusion of PCs.

## Conclusion

We sought to answer whether the standard adjustment for population stratification, through genetic PCs, in GWAS and in polygenic prediction models attenuates direct genetic effect signals.

Our analysis reveals that PCs may tag direct genetic effects, albeit to a modest extent, for a few traits. Namely, top PCs have significant effects on differences between siblings in height and educational attainment. While previous studies have usually not found quadratic effects at the individual allelic level (especially in the case of educational attainment where a dominance GWAS was conducted [Okbay et al. 2022]), we find some evidence of quadratic effects when using PCs (especially for PC 2) for education, height, birthweight, blood pressure and number of drinks per week. For depression and the number of children ever born,we also found significant PC-by-PC interaction effects.

Are estimates of direct genetic effects being attenuated by adjusting for PCs? Our analysis demonstrates that the inclusion of PCs in PGI models does not substantially attenuate the estimates of genetic effects in within family models. This was true both for PGIs based on published GWASs that varied in the homogeneity of cohorts and PC adjustments, and for PGIs based on GWASs that we conducted in the unrelated UKB sample which included the same white British PCs across all phenotypes. That we do not see any changes in the PGI estimatess may, in part, be due to the fact that all GWASs considered here, adjusted for PCs to some extent.

Many of the significant effects of PCs found here are for height and educational attainment. Why might this be the case? One explanation is that both of these traits have been shown to be highly heritable and we may be more powered to detect effects. It could also be due to drift, directional, or stabilizing selection. As Harpak and Przewoski (2021) show, under a wide range of evolutionary models, two genetic groups (however defined) are expected to diverge somewhat in polygenic effects. The within-family estimates of PCs might also be inflated by linkage of alleles on the same chromosome brought about by assortative mating (Veller & Coop 2023). Indeed, height and educational attainment are among the phenotypes that show some of the highest levels of assortative mating, both phenotypically and genotypically (Yengo et al. 2018; Conley et al. 2016).^7^ Assortative mating and selection, have in turn, been shown to be linked (Nishi et al 2020).

Howe et al. (2022) conducted within family GWASs for 25 phenotypes and found that height, educational attainment, and fertility behaviors see the most attenuation in their direct genetic effect estimates (compared to results from traditional between-family GWAS models; see Figure 1 in Howe et al. 2022). This attenuation, especially for more socially mediated phenotypes like educational attainment, is likely due to dynastic confounding of the genetic effects (e.g., generational advantage being passed down alongside genes) (Nivard et al. 2024), and to a lesser extent indirect genetic effects (i.e., environmentally-mediated effects of the shared genes of the proband’s relatives [Kong et al. 2018; Trejo & Domingue 2018]). However, the attenuation could also reflect genetic differences across subpopulations that result from selection, drift, or assortative mating. If rates of mating across these subpopulations are low, then within-family designs will not be able to detect such effects. Thus, while within-family GWAS has long been preferable to GWAS with PC-controls among unrelated individual for the prevention of false positives resulting from population structure, our results suggest that both approaches may sometimes result in false negatives (albeit of relatively modest magnitude). That being said, there is good reason to adjust for confounding, even at the cost of losing a modest degree of causal signal. Future studies can aim to tease apart genetic from non-genetic contributions to population structure.

## Acknowledgements

We thank Evelina Akimova, Aysu Okbay and Patrick Turley for helpful comments on the manuscript. Previous versions of the paper were presented to, and benefited from comments by, members of the Princeton Biosociology Lab, the New York Genome Center, and the 2023 Integrating Genetics and Social Science Conference. A.H was supported by NIH grant R35GM151108.

## Footnotes

1 In genomics, the term ‘direct genetic effect’ is used to describe the causal effect of an organism’s own DNA on that same organism’s traits (whereas the term ‘indirect genetic effect’ refers to the causal effect of an organism’s own DNA on a different organism’s traits). However, ‘direct effects’ and ‘indirect effects’ often carry a different meaning in statistics and the social sciences, where a causal effect is direct if it does not operate through a given set of mediating pathways. In this paper, we utilize the genomic conceptualization of the term. While our within-family design allows us to separate direct genetic effects from confounding indirect genetic effects (e.g., genetic nurture), it does not allow us to separate biologically proximal genetic effects from environmentally mediated genetic effects. Indeed, the precise pathways through which the PCs and PGIs we study have causal effects on traits remains largely unknown and may include complex causal chains involving social and physical environmental features.

2 That said, to the extent that the environment is a mediator of the effects of genetic population structure, it remains (under the potential outcomes framework [Rubin 2005]) an important mechanism for causal genetic effects. For example, if we found that the first PC constructed in the UK Biobank sample predicted smoking behavior within sibling models, it could still be the case that environmental pathways lead the sibling with phenotypes commonly associated with, say, being from the North, or looking more Scottish, to drive effects on smoking, even within families. Such a dynamic might obtain if for example, Scottish-associated phenotypes leads one sibling to have more Scottish friends who also happen to have higher smoking rates in England (https://digital.nhs.uk/data-and-information/publications/statistical/health-survey-england-additional-analyses/ethnicity-and-health-2011-2019-experimental-statistics/cigarette-smoking). A similar argument can be made about skin color and 1000 genomes PCs. Skin tone differences may lead to more stress due to discrimination which might were skin-toned based discrimination that could exact a great toll on darker-skinned siblings— which evidence shows there, in fact, is in the U.S., Brazil and South Africa, at least (see for example, Laidley et al 2019).

3 Veller, Przeworski and Coop (2023) show that even for within family models, genome-wide measures (such as PGIs or, in this case, PCs) cannot be interpreted as the average treatment effect (ATE) since genetic effects are likely heterogeneous by environments and genotypes are non-randomly distributed across those environments (specifically, families differ in their heterozygosity rates across loci). Here, however, we are not interested in identifying the ATE estimates of each PC on a phenotype, but instead whether PCs in general capture direct genetic effects.

4 This low statistical power results from a combination of two factors: 1) the reduced amount of genetic variation within families and 2) the relative paucity of data of dyadic genetic data within families (i.e., parent-child pairs and/or sibling pairs). For instance, though the UK Biobank has roughly 500,000 genotyped individuals, it has only about 22,000 sibling pairs (and even fewer parent-child pairs).

5 All variables are standardized before regressions. Because height is strongly influenced by sex, we standardize height separately for men and women.

6 This is because each set includes the previous 1:*n*th PCs as well, and a comparison with the base model would not allow us to identify the effect of the four focal PCs.

7 Examining the performance of PGI constructed using population-level GWASs, Fletcher et al (2024) show that the R^2^ of these models may be biased upward or downward depending on the environmental correlation of the siblings. This may be consequential for our analyses, but there is no way to empirically test the presence or absence of a systematic environmental correlation among siblings. In addition, this kind of bias is unlikely to only show up for educational attainment and height and no other phenotypes. Fletcher et al (2024) also discuss the bias that could result from the relationship between direct and indirect genetic effects, but that bias is induced by using PGIs trained on population GWASs, which are not the primary focus of the present study.

