## Supplementary Information for "Does standard adjustment for genomic population structure capture direct genetic effects?"

#### Table S1: $R^2$ across different model types and phenotypes

| | *Age + Sex + Array* | *40 Proband PCs + controls* | *40 Parental PCs + controls* | *Full Model* |
|---|---|---|---|---|
| | (1) | (2) | (3) | (4) |
| **BMI** | 0.010 | 0.012 | 0.013 | 0.014 |
| **Height*** | 0.035 | 0.046 | 0.047 | 0.049 |
| **Educational Attainment** | 0.020 | 0.025 | 0.025 | 0.027 |
| **Num Children Ever Born** | 0.029 | 0.031 | 0.031 | 0.033 |
| **Current Smoking** | 0.012 | 0.015 | 0.015 | 0.017 |
| **Drinks per Week** | 0.039 | 0.041 | 0.041 | 0.042 |
| **Diastolic BP** | 0.033 | 0.034 | 0.034 | 0.036 |
| **Depression** | 0.026 | 0.027 | 0.028 | 0.029 |
| **Birthweight** | 0.021 | 0.023 | 0.023 | 0.025 |

$R^2$ across different model specifications is reported. The rows correspond to the phenotypes (the outcomes in the regressions), and the columns correspond to the variables included in the model. Controls refer to age, sex, and array. Full Model includes age, sex, array, 40 parental PCs and 40 proband PCs.
* Height was standardized for males and females separately.

#### Table S2: Percent change in $R^2$ when adding proband PCs to within-family models

| Phenotype | PC set | Incremental R2 (%) | BH corrected p-value |
|---|---|---|---|
| BMI | PCs 1-4 | 0.9702 | 0.5888 |
| BMI | PCs 5-8 | 2.8369 | 0.1199 |
| BMI | PCs 9-12 | 1.0881 | 0.5513 |
| BMI | PCs 13-16 | 1.9025 | 0.2925 |
| BMI | PCs 17-20 | 0.0844 | 0.9987 |
| BMI | PCs 21-24 | 1.1533 | 0.5227 |
| BMI | PCs 25-28 | 1.1404 | 0.5227 |
| BMI | PCs 29-32 | 0.5155 | 0.8139 |
| BMI | PCs 33-36 | 0.8271 | 0.6053 |
| BMI | PCs 37-40 | 1.1873 | 0.5227 |
| Height | PCs 1-4 | 3.0528 | 0.0000 |
| Height | PCs 5-8 | 0.3718 | 0.4664 |
| Height | PCs 9-12 | 0.5624 | 0.2577 |
| Height | PCs 13-16 | 0.4580 | 0.3227 |
| Height | PCs 17-20 | 0.2552 | 0.5697 |
| Height | PCs 21-24 | 0.3912 | 0.4552 |
| Height | PCs 25-28 | 0.7412 | 0.1049 |
| Height | PCs 29-32 | 0.1063 | 0.8974 |

| | | | |
|---|---|---|---|
| Height | PCs 33-36 | 0.2589 | 0.5697 |
| Height | PCs 37-40 | 0.0672 | 0.9750 |
| Educational Attainment | PCs 1-4 | 2.3508 | 0.0052 |
| Educational Attainment | PCs 5-8 | 0.8735 | 0.3238 |
| Educational Attainment | PCs 9-12 | 1.0397 | 0.2708 |
| Educational Attainment | PCs 13-16 | 0.2917 | 0.7698 |
| Educational Attainment | PCs 17-20 | 0.6875 | 0.4664 |
| Educational Attainment | PCs 21-24 | 1.2401 | 0.1612 |
| Educational Attainment | PCs 25-28 | 0.2404 | 0.8451 |
| Educational Attainment | PCs 29-32 | 0.3813 | 0.6569 |
| Educational Attainment | PCs 33-36 | 0.1253 | 0.9750 |
| Educational Attainment | PCs 37-40 | 0.1933 | 0.9007 |
| Number of Children Ever Born | PCs 1-4 | 0.3289 | 0.6489 |
| Number of Children Ever Born | PCs 5-8 | 1.7976 | 0.0052 |
| Number of Children Ever Born | PCs 9-12 | 0.3574 | 0.5888 |
| Number of Children Ever Born | PCs 13-16 | 0.5329 | 0.4962 |
| Number of Children Ever Born | PCs 17-20 | 0.7482 | 0.2925 |
| Number of Children Ever Born | PCs 21-24 | 0.1179 | 0.9643 |
| Number of Children Ever Born | PCs 25-28 | 0.4278 | 0.5513 |
| Number of Children Ever Born | PCs 29-32 | 0.4611 | 0.5227 |
| Number of Children Ever Born | PCs 33-36 | 0.0093 | 0.9987 |
| Number of Children Ever Born | PCs 37-40 | 0.0430 | 0.9987 |
| Cigarettes Per Day | PCs 1-4 | 1.3899 | 0.3686 |
| Cigarettes Per Day | PCs 5-8 | 0.6036 | 0.7005 |
| Cigarettes Per Day | PCs 9-12 | 0.1723 | 0.9750 |
| Cigarettes Per Day | PCs 13-16 | 1.5719 | 0.2925 |
| Cigarettes Per Day | PCs 17-20 | 0.5769 | 0.7018 |
| Cigarettes Per Day | PCs 21-24 | 1.7985 | 0.2458 |
| Cigarettes Per Day | PCs 25-28 | 1.1435 | 0.4664 |
| Cigarettes Per Day | PCs 29-32 | 0.5661 | 0.7005 |
| Cigarettes Per Day | PCs 33-36 | 0.8089 | 0.5697 |
| Cigarettes Per Day | PCs 37-40 | 0.1912 | 0.9750 |
| Drinks Per Week | PCs 1-4 | 0.5790 | 0.2925 |
| Drinks Per Week | PCs 5-8 | 0.3364 | 0.5442 |
| Drinks Per Week | PCs 9-12 | 0.3175 | 0.5624 |
| Drinks Per Week | PCs 13-16 | 0.2055 | 0.7158 |
| Drinks Per Week | PCs 17-20 | 0.0639 | 0.9750 |
| Drinks Per Week | PCs 21-24 | 0.2749 | 0.5888 |
| Drinks Per Week | PCs 25-28 | 0.0062 | 0.9987 |
| Drinks Per Week | PCs 29-32 | 0.2246 | 0.6865 |

| | | | |
|---|---|---|---|
| Drinks Per Week | PCs 33-36 | 0.4412 | 0.4626 |
| Drinks Per Week | PCs 37-40 | 0.0933 | 0.9643 |
| Diastolic BP | PCs 1-4 | 0.3891 | 0.5697 |
| Diastolic BP | PCs 5-8 | 0.0521 | 0.9987 |
| Diastolic BP | PCs 9-12 | 0.9565 | 0.1697 |
| Diastolic BP | PCs 13-16 | 0.4706 | 0.5227 |
| Diastolic BP | PCs 17-20 | 0.5420 | 0.4664 |
| Diastolic BP | PCs 21-24 | 0.3596 | 0.5888 |
| Diastolic BP | PCs 25-28 | 0.1223 | 0.9643 |
| Diastolic BP | PCs 29-32 | 0.4925 | 0.5227 |
| Diastolic BP | PCs 33-36 | 0.3422 | 0.5888 |
| Diastolic BP | PCs 37-40 | 0.2367 | 0.7698 |
| Seen Doc/GP for Depression | PCs 1-4 | 0.4337 | 0.5888 |
| Seen Doc/GP for Depression | PCs 5-8 | 0.4678 | 0.5697 |
| Seen Doc/GP for Depression | PCs 9-12 | 0.2172 | 0.8590 |
| Seen Doc/GP for Depression | PCs 13-16 | 0.3828 | 0.6355 |
| Seen Doc/GP for Depression | PCs 17-20 | 0.5670 | 0.5227 |
| Seen Doc/GP for Depression | PCs 21-24 | 0.1938 | 0.8974 |
| Seen Doc/GP for Depression | PCs 25-28 | 0.8476 | 0.2944 |
| Seen Doc/GP for Depression | PCs 29-32 | 0.8262 | 0.2956 |
| Seen Doc/GP for Depression | PCs 33-36 | 0.5190 | 0.5287 |
| Seen Doc/GP for Depression | PCs 37-40 | 0.2617 | 0.7698 |
| Birthweight | PCs 1-4 | 1.0348 | 0.5442 |
| Birthweight | PCs 5-8 | 0.5545 | 0.7698 |
| Birthweight | PCs 9-12 | 0.0676 | 0.9987 |
| Birthweight | PCs 13-16 | 0.4497 | 0.8451 |
| Birthweight | PCs 17-20 | 1.1046 | 0.5227 |
| Birthweight | PCs 21-24 | 0.5289 | 0.7698 |
| Birthweight | PCs 25-28 | 0.1772 | 0.9750 |
| Birthweight | PCs 29-32 | 3.0773 | 0.0368 |
| Birthweight | PCs 33-36 | 0.2046 | 0.9750 |
| Birthweight | PCs 37-40 | 0.2588 | 0.9643 |

Percent change in $R^2$, when adding proband PCs compared to the base model (age + sex + array + parental PCs), for each phenotype are reported. PCs are added in bundles of 4, cumulatively. They portray the effect of PC n:n+4 while controlling for PCs 1:n. BH-corrected p-values are reported for model comparisons using F-tests.

**Table S3: Percent change in $R^2$ in bivariate interaction and quadratic models of PCs 1-8 and PGIs**

| Phenotype | PC set | Incremental R2 (%) | BH corrected p-value |
|---|---|---|---|
| Educational Attainment | PC 2 x PC 2 | 3.75058 | 0.00002 |
| Height | PC 3 x PC 4 | 1.90121 | 0.00003 |
| Height | PC 2 x PC 2 | 1.73312 | 0.00011 |
| Educational Attainment | PC 1 x PC 2 | 2.60395 | 0.00047 |
| Height | PC 2 x PC 4 | 1.37261 | 0.00094 |
| Drinks Per Week | PC 1 x PC 2 | 1.15856 | 0.00119 |
| Educational Attainment | PC 1 x PC 5 | 2.04588 | 0.00210 |
| Seen Doc/GP for Depression | PC 2 x PC 4 | 1.53148 | 0.00289 |
| Birthweight | PC 4 x PC 4 | 3.19564 | 0.00398 |
| Educational Attainment | PC 5 x PGI | 0.64857 | 0.00590 |
| Number of Children Ever Born | PC 1 x PC 3 | 1.06156 | 0.01488 |
| Educational Attainment | PC 3 x PC 3 | 1.42539 | 0.01834 |
| Diastolic BP | PC 2 x PC 2 | 0.95702 | 0.01834 |
| Drinks Per Week | PC 2 x PC 2 | 0.73151 | 0.01862 |
| Diastolic BP | PC 1 x PC 5 | 0.91079 | 0.02159 |
| Height | PC 3 x PC 3 | 0.62962 | 0.06061 |
| Number of Children Ever Born | PC 1 x PC 2 | 0.77328 | 0.06061 |
| Educational Attainment | PC 3 x PC 4 | 1.01337 | 0.06624 |
| BMI | PC 3 x PC 3 | 2.19423 | 0.07326 |
| Educational Attainment | PC 2 x PC 3 | 1.00967 | 0.07326 |
| Number of Children Ever Born | PC 2 x PC 4 | 0.72631 | 0.07326 |
| Drinks Per Week | PC 1 x PC 3 | 0.51171 | 0.07952 |
| Height | PC 5 x PC 5 | 0.54042 | 0.09271 |
| Drinks Per Week | PC 3 x PC 3 | 0.48519 | 0.09271 |
| Cigarettes Per Day | PC 1 x PC 1 | 1.43463 | 0.10272 |
| Diastolic BP | PC 2 x PC 3 | 0.55396 | 0.13299 |
| Cigarettes Per Day | PC 4 x PGI | 1.26464 | 0.13659 |
| Height | PC 4 x PC 4 | 0.47997 | 0.13896 |
| Educational Attainment | PC 4 x PC 4 | 0.75177 | 0.16458 |
| Cigarettes Per Day | PC 1 x PC 8 | 1.17463 | 0.16833 |
| BMI | PC 1 x PGI | 0.36599 | 0.17028 |
| Number of Children Ever Born | PC 2 x PC 7 | 0.52434 | 0.17668 |
| Cigarettes Per Day | PC 3 x PC 3 | 1.21962 | 0.17868 |
| Height | PC 2 x PC 3 | 0.39939 | 0.18581 |
| Number of Children Ever Born | PC 5 x PC 5 | 0.47968 | 0.19247 |
| Number of Children Ever Born | PC 2 x PC 2 | 0.49198 | 0.19425 |

| | | | |
|---|---|---|---|
| Drinks Per Week | PC 4 x PC 4 | 0.34955 | 0.19425 |
| Educational Attainment | PC 5 x PC 6 | 0.65763 | 0.19728 |
| Educational Attainment | PC 6 x PC 6 | 0.68556 | 0.19728 |
| Birthweight | PC 1 x PC 5 | 1.11519 | 0.19728 |
| BMI | PC 3 x PGI | 0.32047 | 0.20304 |
| Number of Children Ever Born | PC 1 x PC 1 | 0.44111 | 0.21256 |
| Cigarettes Per Day | PC 6 x PC 8 | 1.03489 | 0.21256 |
| Cigarettes Per Day | PC 2 x PC 4 | 1.04446 | 0.22051 |
| Seen Doc/GP for Depression | PC 2 x PC 3 | 0.47727 | 0.22514 |
| BMI | PC 2 x PC 4 | 1.18770 | 0.27318 |
| Height | PC 1 x PC 5 | 0.26084 | 0.27318 |
| Drinks Per Week | PC 3 x PC 4 | 0.27947 | 0.27613 |
| Birthweight | PC 3 x PC 7 | 0.93517 | 0.27613 |
| Educational Attainment | PC 4 x PC 5 | 0.50532 | 0.28728 |
| Cigarettes Per Day | PC 1 x PC 6 | 0.82504 | 0.28728 |
| Height | PC 2 x PC 5 | 0.29707 | 0.29499 |
| Number of Children Ever Born | PC 2 x PC 3 | 0.37098 | 0.29499 |
| BMI | PC 5 x PC 8 | 1.05532 | 0.30280 |
| Educational Attainment | PC 6 x PC 7 | 0.51851 | 0.30280 |
| Seen Doc/GP for Depression | PC 3 x PC 3 | 0.38140 | 0.32648 |
| Educational Attainment | PC 3 x PC 5 | 0.44747 | 0.34684 |
| Cigarettes Per Day | PC 3 x PC 7 | 0.78630 | 0.34684 |
| Diastolic BP | PC 1 x PC 7 | 0.30103 | 0.35242 |
| Birthweight | PC 2 x PC 6 | 0.74251 | 0.35242 |
| Birthweight | PC 4 x PC 5 | 0.78016 | 0.35242 |
| Drinks Per Week | PC 1 x PC 1 | 0.22907 | 0.36736 |
| Educational Attainment | PC 1 x PC 3 | 0.41611 | 0.39664 |
| Diastolic BP | PC 8 x PC 8 | 0.27511 | 0.39950 |
| Seen Doc/GP for Depression | PC 5 x PC 8 | 0.32565 | 0.39950 |
| BMI | PC 2 x PC 2 | 0.88644 | 0.41023 |
| Number of Children Ever Born | PC 1 x PC 4 | 0.27594 | 0.41812 |
| Birthweight | PC 7 x PC 8 | 0.67479 | 0.41812 |
| Height | PC 7 x PC 8 | 0.22724 | 0.42823 |
| Number of Children Ever Born | PC 8 x PC 8 | 0.27574 | 0.42980 |
| Educational Attainment | PC 1 x PC 7 | 0.38963 | 0.43156 |
| Educational Attainment | PC 1 x PC 8 | 0.38191 | 0.44019 |
| Height | PC 4 x PC 8 | 0.20912 | 0.45596 |
| Educational Attainment | PC 7 x PC 8 | 0.36581 | 0.45596 |
| Diastolic BP | PC 1 x PC 4 | 0.24007 | 0.45596 |
| Birthweight | PC 1 x PGI | 0.39196 | 0.45596 |

| | | | |
|---|---|---|---|
| BMI | PC 1 x PC 7 | 0.70202 | 0.45837 |
| Number of Children Ever Born | PC 4 x PC 6 | 0.24987 | 0.45837 |
| Drinks Per Week | PC 7 x PC 7 | 0.17593 | 0.47516 |
| Diastolic BP | PC 6 x PC 8 | 0.22579 | 0.47516 |
| Number of Children Ever Born | PC 3 x PC 8 | 0.23808 | 0.48188 |
| Drinks Per Week | PC 2 x PGI | 0.15539 | 0.49478 |
| Diastolic BP | PC 3 x PC 6 | 0.21505 | 0.49478 |
| Height | PC 8 x PGI | 0.05978 | 0.49682 |
| Educational Attainment | PC 3 x PC 7 | 0.31068 | 0.49682 |
| Cigarettes Per Day | PC 4 x PC 4 | 0.54175 | 0.49682 |
| Birthweight | PC 1 x PC 4 | 0.52936 | 0.49682 |
| BMI | PC 3 x PC 6 | 0.63465 | 0.51721 |
| BMI | PC 1 x PC 3 | 0.56343 | 0.54475 |
| Number of Children Ever Born | PC 3 x PC 3 | 0.20652 | 0.54571 |
| Birthweight | PC 4 x PC 6 | 0.48300 | 0.55533 |
| Diastolic BP | PC 6 x PC 6 | 0.18657 | 0.55978 |
| Height | PC 3 x PC 5 | 0.14750 | 0.57419 |
| Height | PC 2 x PGI | 0.05113 | 0.57419 |
| Educational Attainment | PC 5 x PC 7 | 0.26138 | 0.57419 |
| Drinks Per Week | PC 4 x PGI | 0.13002 | 0.57419 |
| Diastolic BP | PC 5 x PC 8 | 0.17494 | 0.57419 |
| Birthweight | PC 5 x PC 7 | 0.44714 | 0.57419 |
| BMI | PC 3 x PC 4 | 0.53186 | 0.58152 |
| Number of Children Ever Born | PC 1 x PC 5 | 0.17367 | 0.58152 |
| Height | PC 1 x PC 6 | 0.11492 | 0.59667 |
| Number of Children Ever Born | PC 5 x PC 6 | 0.16497 | 0.59667 |
| Number of Children Ever Born | PC 3 x PGI | 0.16189 | 0.59667 |
| Number of Children Ever Born | PC 4 x PGI | 0.16106 | 0.59667 |
| Cigarettes Per Day | PC 5 x PC 8 | 0.38570 | 0.59667 |
| Drinks Per Week | PC 1 x PC 5 | 0.12552 | 0.59667 |
| Drinks Per Week | PC 6 x PGI | 0.11503 | 0.59667 |
| Diastolic BP | PC 7 x PC 8 | 0.15849 | 0.59667 |
| Educational Attainment | PC 4 x PC 6 | 0.22889 | 0.60129 |
| Seen Doc/GP for Depression | PC 5 x PC 5 | 0.18200 | 0.60322 |
| Number of Children Ever Born | PC 6 x PC 8 | 0.16182 | 0.61952 |
| BMI | PC 2 x PC 3 | 0.46511 | 0.63423 |
| Height | PC 6 x PC 8 | 0.12762 | 0.63423 |
| BMI | PC 1 x PC 5 | 0.39765 | 0.64958 |
| BMI | PC 4 x PC 7 | 0.44288 | 0.64958 |
| BMI | PC 4 x PC 4 | 0.44900 | 0.64958 |

| | | | |
|---|---|---|---|
| Number of Children Ever Born | PC 8 x PGI | 0.13897 | 0.64958 |
| Cigarettes Per Day | PC 4 x PC 5 | 0.33914 | 0.64958 |
| Drinks Per Week | PC 6 x PC 6 | 0.10689 | 0.64958 |
| Seen Doc/GP for Depression | PC 3 x PC 6 | 0.16020 | 0.64958 |
| Cigarettes Per Day | PC 3 x PC 8 | 0.32746 | 0.65504 |
| Cigarettes Per Day | PC 2 x PGI | 0.30801 | 0.65504 |
| BMI | PC 7 x PGI | 0.09480 | 0.67030 |
| Height | PC 1 x PC 4 | 0.08930 | 0.67030 |
| Height | PC 4 x PGI | 0.03639 | 0.67030 |
| Birthweight | PC 3 x PC 4 | 0.32261 | 0.67030 |
| Educational Attainment | PC 6 x PC 8 | 0.18929 | 0.67109 |
| Number of Children Ever Born | PC 1 x PC 7 | 0.12969 | 0.67109 |
| Cigarettes Per Day | PC 1 x PC 2 | 0.29378 | 0.67109 |
| Diastolic BP | PC 4 x PC 5 | 0.12138 | 0.67422 |
| Number of Children Ever Born | PC 1 x PC 6 | 0.12290 | 0.69120 |
| Diastolic BP | PC 4 x PC 4 | 0.11682 | 0.69120 |
| Drinks Per Week | PC 1 x PC 8 | 0.09031 | 0.69341 |
| Cigarettes Per Day | PC 5 x PC 7 | 0.27841 | 0.69982 |
| Cigarettes Per Day | PC 6 x PGI | 0.25278 | 0.69982 |
| Drinks Per Week | PC 1 x PC 4 | 0.08704 | 0.69982 |
| Birthweight | PC 2 x PGI | 0.17773 | 0.69982 |
| Number of Children Ever Born | PC 5 x PC 8 | 0.11309 | 0.70714 |
| Number of Children Ever Born | PC 5 x PGI | 0.10417 | 0.71010 |
| Birthweight | PC 3 x PC 5 | 0.27367 | 0.71314 |
| Cigarettes Per Day | PC 4 x PC 8 | 0.26131 | 0.71828 |
| Drinks Per Week | PC 2 x PC 7 | 0.08018 | 0.72355 |
| Height | PC 7 x PGI | 0.02833 | 0.72736 |
| Height | PC 1 x PC 1 | 0.07211 | 0.72736 |
| Educational Attainment | PC 2 x PGI | 0.05537 | 0.72736 |
| Drinks Per Week | PC 2 x PC 3 | 0.07705 | 0.72736 |
| Seen Doc/GP for Depression | PC 2 x PC 2 | 0.11865 | 0.72736 |
| Educational Attainment | PC 7 x PGI | 0.05354 | 0.73065 |
| Educational Attainment | PC 5 x PC 5 | 0.14464 | 0.73065 |
| Diastolic BP | PC 1 x PGI | 0.09675 | 0.73065 |
| Birthweight | PC 5 x PC 6 | 0.24650 | 0.73065 |
| Drinks Per Week | PC 5 x PC 7 | 0.07206 | 0.73677 |
| Diastolic BP | PC 4 x PC 6 | 0.09289 | 0.73677 |
| Drinks Per Week | PC 6 x PC 7 | 0.07104 | 0.74085 |
| Diastolic BP | PC 6 x PC 7 | 0.09132 | 0.74085 |
| Cigarettes Per Day | PC 3 x PGI | 0.20634 | 0.74180 |

| | | | |
|---|---|---|---|
| BMI | PC 4 x PC 5 | 0.27052 | 0.75509 |
| Height | PC 5 x PC 7 | 0.07515 | 0.75509 |
| BMI | PC 1 x PC 6 | 0.24688 | 0.76266 |
| BMI | PC 3 x PC 7 | 0.26642 | 0.76266 |
| Number of Children Ever Born | PC 2 x PGI | 0.08304 | 0.76266 |
| Drinks Per Week | PC 3 x PC 6 | 0.06539 | 0.76266 |
| Seen Doc/GP for Depression | PC 3 x PC 5 | 0.09594 | 0.76859 |
| Educational Attainment | PC 1 x PGI | 0.04520 | 0.76956 |
| Diastolic BP | PC 3 x PC 5 | 0.07851 | 0.77836 |
| BMI | PC 2 x PC 6 | 0.24687 | 0.78231 |
| BMI | PC 8 x PC 8 | 0.24981 | 0.78231 |
| Birthweight | PC 5 x PGI | 0.12502 | 0.78231 |
| BMI | PC 7 x PC 8 | 0.24236 | 0.78366 |
| Number of Children Ever Born | PC 1 x PC 8 | 0.07813 | 0.78366 |
| Drinks Per Week | PC 2 x PC 4 | 0.05654 | 0.78366 |
| Diastolic BP | PC 2 x PC 7 | 0.07400 | 0.78366 |
| Number of Children Ever Born | PC 3 x PC 6 | 0.07764 | 0.79022 |
| Height | PC 1 x PC 8 | 0.04934 | 0.79060 |
| Height | PC 2 x PC 6 | 0.06160 | 0.79060 |
| Height | PC 3 x PGI | 0.02001 | 0.79060 |
| Seen Doc/GP for Depression | PC 1 x PC 1 | 0.08180 | 0.79060 |
| Birthweight | PC 6 x PC 7 | 0.17966 | 0.79060 |
| Birthweight | PC 7 x PC 7 | 0.17482 | 0.79354 |
| BMI | PC 1 x PC 1 | 0.19986 | 0.79430 |
| Educational Attainment | PC 2 x PC 5 | 0.09914 | 0.79430 |
| Drinks Per Week | PC 2 x PC 6 | 0.05083 | 0.79430 |
| Drinks Per Week | PC 5 x PGI | 0.04743 | 0.79430 |
| Seen Doc/GP for Depression | PC 1 x PC 7 | 0.07714 | 0.79430 |
| Educational Attainment | PC 4 x PC 7 | 0.09304 | 0.80262 |
| Number of Children Ever Born | PC 6 x PGI | 0.06365 | 0.80262 |
| Seen Doc/GP for Depression | PC 8 x PC 8 | 0.07204 | 0.81701 |
| Birthweight | PC 2 x PC 4 | 0.14916 | 0.81701 |
| Birthweight | PC 2 x PC 3 | 0.14751 | 0.81869 |
| Birthweight | PC 1 x PC 6 | 0.14834 | 0.81967 |
| Birthweight | PC 3 x PGI | 0.09576 | 0.81967 |
| Drinks Per Week | PC 3 x PC 7 | 0.04251 | 0.83228 |
| Seen Doc/GP for Depression | PC 1 x PC 6 | 0.06570 | 0.83228 |
| Seen Doc/GP for Depression | PC 5 x PC 6 | 0.06545 | 0.83228 |
| Seen Doc/GP for Depression | PC 4 x PC 4 | 0.06420 | 0.83228 |
| Birthweight | PC 2 x PC 2 | 0.13808 | 0.83228 |

| | | | |
|---|---|---|---|
| Height | PC 2 x PC 8 | 0.04746 | 0.83306 |
| Educational Attainment | PC 3 x PC 8 | 0.07946 | 0.83455 |
| Diastolic BP | PC 1 x PC 6 | 0.05289 | 0.83575 |
| Cigarettes Per Day | PC 2 x PC 7 | 0.13174 | 0.83971 |
| Diastolic BP | PC 1 x PC 3 | 0.05141 | 0.83971 |
| Birthweight | PC 4 x PC 8 | 0.12882 | 0.84651 |
| Number of Children Ever Born | PC 6 x PC 6 | 0.05297 | 0.85132 |
| BMI | PC 8 x PGI | 0.03513 | 0.85486 |
| Cigarettes Per Day | PC 7 x PC 8 | 0.11870 | 0.85486 |
| Drinks Per Week | PC 4 x PC 6 | 0.03621 | 0.85486 |
| Diastolic BP | PC 2 x PC 4 | 0.04710 | 0.85486 |
| BMI | PC 3 x PC 5 | 0.14255 | 0.85575 |
| Cigarettes Per Day | PC 5 x PGI | 0.10255 | 0.85575 |
| Seen Doc/GP for Depression | PC 3 x PC 7 | 0.05384 | 0.85575 |
| BMI | PC 1 x PC 4 | 0.12642 | 0.85754 |
| BMI | PC 6 x PC 7 | 0.12856 | 0.85754 |
| BMI | PC 5 x PC 5 | 0.12478 | 0.85754 |
| Educational Attainment | PC 2 x PC 6 | 0.06838 | 0.85754 |
| Educational Attainment | PC 3 x PC 6 | 0.05967 | 0.85754 |
| Cigarettes Per Day | PC 1 x PC 5 | 0.09394 | 0.85754 |
| Drinks Per Week | PC 3 x PC 8 | 0.02961 | 0.85754 |
| Diastolic BP | PC 2 x PC 5 | 0.04287 | 0.85754 |
| Diastolic BP | PC 3 x PC 8 | 0.04327 | 0.85754 |
| Seen Doc/GP for Depression | PC 1 x PC 2 | 0.04859 | 0.85754 |
| Seen Doc/GP for Depression | PC 1 x PC 5 | 0.05174 | 0.85754 |
| Seen Doc/GP for Depression | PC 2 x PC 7 | 0.04570 | 0.85754 |
| Seen Doc/GP for Depression | PC 3 x PC 4 | 0.04672 | 0.85754 |
| Seen Doc/GP for Depression | PC 6 x PC 8 | 0.04627 | 0.85754 |
| Seen Doc/GP for Depression | PC 8 x PGI | 0.04901 | 0.85754 |
| Birthweight | PC 2 x PC 5 | 0.09398 | 0.85754 |
| Birthweight | PC 2 x PC 7 | 0.09500 | 0.85754 |
| Birthweight | PC 8 x PC 8 | 0.10507 | 0.85754 |
| BMI | PC 6 x PGI | 0.02747 | 0.86317 |
| Diastolic BP | PC 5 x PC 5 | 0.03680 | 0.86317 |
| BMI | PC 5 x PC 7 | 0.10895 | 0.87420 |
| Height | PC 6 x PC 7 | 0.03030 | 0.87420 |
| Height | PC 8 x PC 8 | 0.02845 | 0.87420 |
| Educational Attainment | PC 2 x PC 7 | 0.05159 | 0.87420 |
| Educational Attainment | PC 8 x PGI | 0.01808 | 0.87420 |
| Number of Children Ever Born | PC 7 x PC 7 | 0.03777 | 0.87420 |

| | | | |
|---|---|---|---|
| Cigarettes Per Day | PC 2 x PC 3 | 0.08079 | 0.87420 |
| Cigarettes Per Day | PC 3 x PC 6 | 0.07780 | 0.87420 |
| Diastolic BP | PC 1 x PC 8 | 0.03411 | 0.87420 |
| Seen Doc/GP for Depression | PC 1 x PC 3 | 0.03849 | 0.87420 |
| Seen Doc/GP for Depression | PC 2 x PC 8 | 0.03627 | 0.87420 |
| Birthweight | PC 6 x PC 8 | 0.08129 | 0.87420 |
| Birthweight | PC 3 x PC 3 | 0.07890 | 0.87420 |
| Birthweight | PC 6 x PC 6 | 0.07978 | 0.87420 |
| Drinks Per Week | PC 7 x PGI | 0.02173 | 0.87957 |
| Height | PC 4 x PC 5 | 0.02493 | 0.88024 |
| Educational Attainment | PC 3 x PGI | 0.01566 | 0.88024 |
| Number of Children Ever Born | PC 4 x PC 8 | 0.03097 | 0.88024 |
| Cigarettes Per Day | PC 2 x PC 8 | 0.07161 | 0.88024 |
| Diastolic BP | PC 3 x PC 4 | 0.02802 | 0.88024 |
| Height | PC 4 x PC 7 | 0.02425 | 0.88523 |
| Height | PC 5 x PGI | 0.00728 | 0.88573 |
| Cigarettes Per Day | PC 5 x PC 6 | 0.06325 | 0.88573 |
| Diastolic BP | PC 2 x PC 8 | 0.02680 | 0.88573 |
| Diastolic BP | PC 4 x PC 7 | 0.02645 | 0.88573 |
| Seen Doc/GP for Depression | PC 1 x PGI | 0.03075 | 0.88573 |
| Birthweight | PC 3 x PC 8 | 0.06500 | 0.88573 |
| Birthweight | PC 5 x PC 5 | 0.06507 | 0.88573 |
| BMI | PC 4 x PC 6 | 0.06602 | 0.89409 |
| Height | PC 3 x PC 8 | 0.01818 | 0.89409 |
| Educational Attainment | PC 8 x PC 8 | 0.03455 | 0.89409 |
| Number of Children Ever Born | PC 7 x PGI | 0.02127 | 0.89409 |
| Cigarettes Per Day | PC 3 x PC 4 | 0.05475 | 0.89409 |
| Cigarettes Per Day | PC 7 x PGI | 0.05343 | 0.89409 |
| Drinks Per Week | PC 2 x PC 8 | 0.01699 | 0.89409 |
| Drinks Per Week | PC 3 x PC 5 | 0.01841 | 0.89409 |
| Seen Doc/GP for Depression | PC 1 x PC 8 | 0.02486 | 0.89409 |
| Seen Doc/GP for Depression | PC 4 x PC 8 | 0.02466 | 0.89409 |
| Seen Doc/GP for Depression | PC 6 x PC 7 | 0.02458 | 0.89409 |
| Seen Doc/GP for Depression | PC 6 x PC 6 | 0.02738 | 0.89409 |
| Seen Doc/GP for Depression | PC 7 x PC 7 | 0.02459 | 0.89409 |
| Cigarettes Per Day | PC 8 x PC 8 | 0.05161 | 0.89577 |
| Seen Doc/GP for Depression | PC 3 x PGI | 0.02360 | 0.89795 |
| Number of Children Ever Born | PC 2 x PC 8 | 0.02048 | 0.90063 |
| Diastolic BP | PC 1 x PC 1 | 0.01926 | 0.90063 |
| Seen Doc/GP for Depression | PC 1 x PC 4 | 0.02201 | 0.90063 |

| | | | |
|---|---|---|---|
| Birthweight | PC 1 x PC 8 | 0.04862 | 0.90063 |
| Birthweight | PC 3 x PC 6 | 0.04734 | 0.90234 |
| BMI | PC 2 x PC 7 | 0.05625 | 0.90404 |
| Number of Children Ever Born | PC 2 x PC 6 | 0.01861 | 0.90404 |
| Cigarettes Per Day | PC 1 x PGI | 0.03893 | 0.90404 |
| Diastolic BP | PC 3 x PC 3 | 0.01769 | 0.90404 |
| Seen Doc/GP for Depression | PC 5 x PC 7 | 0.02029 | 0.90404 |
| Drinks Per Week | PC 1 x PC 7 | 0.01279 | 0.90493 |
| Diastolic BP | PC 7 x PGI | 0.01671 | 0.90493 |
| Birthweight | PC 1 x PC 3 | 0.04157 | 0.90493 |
| BMI | PC 1 x PC 2 | 0.04304 | 0.90664 |
| BMI | PC 7 x PC 7 | 0.04793 | 0.90664 |
| Educational Attainment | PC 6 x PGI | 0.00888 | 0.90664 |
| Number of Children Ever Born | PC 5 x PC 7 | 0.01527 | 0.90664 |
| Cigarettes Per Day | PC 6 x PC 7 | 0.03562 | 0.90664 |
| Cigarettes Per Day | PC 8 x PGI | 0.03339 | 0.90664 |
| Drinks Per Week | PC 7 x PC 8 | 0.01221 | 0.90664 |
| Drinks Per Week | PC 1 x PGI | 0.01066 | 0.90664 |
| Seen Doc/GP for Depression | PC 5 x PGI | 0.01686 | 0.90664 |
| Cigarettes Per Day | PC 4 x PC 7 | 0.03516 | 0.90957 |
| Height | PC 6 x PC 6 | 0.01170 | 0.91528 |
| BMI | PC 2 x PC 5 | 0.03863 | 0.91808 |
| BMI | PC 4 x PGI | 0.00756 | 0.91808 |
| BMI | PC 5 x PGI | 0.00758 | 0.91808 |
| Educational Attainment | PC 1 x PC 4 | 0.01708 | 0.91808 |
| Educational Attainment | PC 4 x PC 8 | 0.01717 | 0.91808 |
| Educational Attainment | PC 5 x PC 8 | 0.01814 | 0.91808 |
| Number of Children Ever Born | PC 4 x PC 5 | 0.01224 | 0.91808 |
| Cigarettes Per Day | PC 2 x PC 6 | 0.02584 | 0.91808 |
| Cigarettes Per Day | PC 3 x PC 5 | 0.02619 | 0.91808 |
| Cigarettes Per Day | PC 6 x PC 6 | 0.02652 | 0.91808 |
| Drinks Per Week | PC 4 x PC 7 | 0.00987 | 0.91808 |
| Drinks Per Week | PC 5 x PC 5 | 0.00859 | 0.91808 |
| Seen Doc/GP for Depression | PC 7 x PC 8 | 0.01342 | 0.91808 |
| Birthweight | PC 1 x PC 2 | 0.02592 | 0.91808 |
| Birthweight | PC 7 x PGI | 0.01786 | 0.91808 |
| Height | PC 3 x PC 7 | 0.00758 | 0.92049 |
| Height | PC 5 x PC 6 | 0.00814 | 0.92049 |
| Cigarettes Per Day | PC 1 x PC 7 | 0.02250 | 0.92049 |
| Diastolic BP | PC 2 x PC 6 | 0.00882 | 0.92049 |

| | | | |
|---|---|---|---|
| Seen Doc/GP for Depression | PC 2 x PC 5 | 0.01038 | 0.92049 |
| Seen Doc/GP for Depression | PC 2 x PC 6 | 0.01090 | 0.92049 |
| Seen Doc/GP for Depression | PC 4 x PGI | 0.01034 | 0.92049 |
| Drinks Per Week | PC 4 x PC 8 | 0.00614 | 0.93102 |
| Diastolic BP | PC 5 x PC 7 | 0.00794 | 0.93102 |
| Height | PC 1 x PC 3 | 0.00490 | 0.93385 |
| Height | PC 4 x PC 6 | 0.00638 | 0.93385 |
| Diastolic BP | PC 1 x PC 2 | 0.00744 | 0.93385 |
| Birthweight | PC 4 x PGI | 0.01210 | 0.93385 |
| BMI | PC 4 x PC 8 | 0.02082 | 0.93506 |
| Drinks Per Week | PC 8 x PGI | 0.00498 | 0.93506 |
| Birthweight | PC 1 x PC 7 | 0.01710 | 0.93506 |
| BMI | PC 1 x PC 8 | 0.01887 | 0.93621 |
| BMI | PC 3 x PC 8 | 0.01513 | 0.93621 |
| BMI | PC 2 x PGI | 0.00450 | 0.93621 |
| Height | PC 1 x PC 2 | 0.00393 | 0.93621 |
| Height | PC 6 x PGI | 0.00138 | 0.93621 |
| Height | PC 7 x PC 7 | 0.00412 | 0.93621 |
| Number of Children Ever Born | PC 6 x PC 7 | 0.00506 | 0.93621 |
| Number of Children Ever Born | PC 1 x PGI | 0.00617 | 0.93621 |
| Cigarettes Per Day | PC 5 x PC 5 | 0.01478 | 0.93621 |
| Drinks Per Week | PC 1 x PC 6 | 0.00404 | 0.93621 |
| Diastolic BP | PC 4 x PGI | 0.00473 | 0.93621 |
| Diastolic BP | PC 5 x PGI | 0.00509 | 0.93621 |
| Diastolic BP | PC 8 x PGI | 0.00571 | 0.93621 |
| Seen Doc/GP for Depression | PC 3 x PC 8 | 0.00688 | 0.93621 |
| Seen Doc/GP for Depression | PC 4 x PC 6 | 0.00593 | 0.93621 |
| Number of Children Ever Born | PC 3 x PC 7 | 0.00484 | 0.93680 |
| Number of Children Ever Born | PC 3 x PC 5 | 0.00397 | 0.94481 |
| Drinks Per Week | PC 5 x PC 6 | 0.00305 | 0.94481 |
| Seen Doc/GP for Depression | PC 4 x PC 5 | 0.00450 | 0.94481 |
| Number of Children Ever Born | PC 2 x PC 5 | 0.00348 | 0.94950 |
| Diastolic BP | PC 4 x PC 8 | 0.00333 | 0.94950 |
| Birthweight | PC 1 x PC 1 | 0.00885 | 0.94950 |
| Drinks Per Week | PC 3 x PGI | 0.00223 | 0.95393 |
| BMI | PC 5 x PC 6 | 0.00726 | 0.95918 |
| BMI | PC 6 x PC 8 | 0.00774 | 0.95918 |
| Height | PC 1 x PC 7 | 0.00182 | 0.95918 |
| Height | PC 5 x PC 8 | 0.00168 | 0.95918 |
| Educational Attainment | PC 2 x PC 4 | 0.00314 | 0.95918 |

| | | | |
|---|---|---|---|
| Educational Attainment | PC 1 x PC 1 | 0.00350 | 0.95918 |
| Educational Attainment | PC 7 x PC 7 | 0.00308 | 0.95918 |
| Number of Children Ever Born | PC 4 x PC 7 | 0.00231 | 0.95918 |
| Cigarettes Per Day | PC 1 x PC 3 | 0.00503 | 0.95918 |
| Diastolic BP | PC 3 x PC 7 | 0.00208 | 0.95918 |
| Seen Doc/GP for Depression | PC 6 x PGI | 0.00314 | 0.95918 |
| Seen Doc/GP for Depression | PC 4 x PC 7 | 0.00210 | 0.96185 |
| Birthweight | PC 2 x PC 8 | 0.00423 | 0.96203 |
| Birthweight | PC 8 x PGI | 0.00269 | 0.96203 |
| BMI | PC 2 x PC 8 | 0.00270 | 0.96469 |
| Height | PC 3 x PC 6 | 0.00083 | 0.96469 |
| Height | PC 1 x PGI | 0.00022 | 0.96469 |
| Educational Attainment | PC 2 x PC 8 | 0.00116 | 0.96469 |
| Educational Attainment | PC 4 x PGI | 0.00079 | 0.96469 |
| Number of Children Ever Born | PC 7 x PC 8 | 0.00116 | 0.96469 |
| Cigarettes Per Day | PC 2 x PC 5 | 0.00176 | 0.96469 |
| Cigarettes Per Day | PC 4 x PC 6 | 0.00353 | 0.96469 |
| Cigarettes Per Day | PC 7 x PC 7 | 0.00349 | 0.96469 |
| Drinks Per Week | PC 2 x PC 5 | 0.00067 | 0.96469 |
| Drinks Per Week | PC 4 x PC 5 | 0.00080 | 0.96469 |
| Drinks Per Week | PC 8 x PC 8 | 0.00063 | 0.96469 |
| Diastolic BP | PC 5 x PC 6 | 0.00088 | 0.96469 |
| Diastolic BP | PC 6 x PGI | 0.00099 | 0.96469 |
| Diastolic BP | PC 7 x PC 7 | 0.00071 | 0.96469 |
| Height | PC 2 x PC 7 | 0.00043 | 0.96777 |
| Drinks Per Week | PC 5 x PC 8 | 0.00045 | 0.96777 |
| Seen Doc/GP for Depression | PC 7 x PGI | 0.00063 | 0.96777 |
| Birthweight | PC 5 x PC 8 | 0.00129 | 0.96777 |
| Cigarettes Per Day | PC 1 x PC 4 | 0.00101 | 0.96909 |
| Diastolic BP | PC 2 x PGI | 0.00033 | 0.97222 |
| Birthweight | PC 4 x PC 7 | 0.00081 | 0.97222 |
| Number of Children Ever Born | PC 4 x PC 4 | 0.00019 | 0.98048 |
| Seen Doc/GP for Depression | PC 2 x PGI | 0.00022 | 0.98048 |
| Drinks Per Week | PC 6 x PC 8 | 0.00010 | 0.98328 |
| Birthweight | PC 6 x PGI | 0.00017 | 0.98365 |
| Number of Children Ever Born | PC 3 x PC 4 | 0.00007 | 0.98677 |
| BMI | PC 6 x PC 6 | 0.00011 | 0.98941 |
| Educational Attainment | PC 1 x PC 6 | 0.00005 | 0.98941 |
| Diastolic BP | PC 3 x PGI | 0.00002 | 0.98946 |
| Cigarettes Per Day | PC 2 x PC 2 | 0.00001 | 0.99542 |

Percent change in $R^2$ compared to the base model are reported for the different interaction models for each phenotype. In addition to proband controls (sex, age, and array), base models include the relevant mid-parental PC or PGI, as well as an additive term for the proband PC or PGI. The models are not cumulative (i.e. prior PCs are not adjusted for). BH-corrected p-values are reported for model comparisons using F-tests.

#### Examining the change in PGI estimates when adjusting for the direct effects of PCs

Having established a modest direct effect of PCs on select phenotypes, we examine whether including PCs as controls in within-family PGI prediction models, leads to a significant change in the estimate of the PGI. Table S3 reports p-values from the Clogg, Petkova, and Haritou (1995) tests comparing PGI coefficients before and after adding PCs. None of the comparisons are statistically significant, which show that while PCs have a small direct effect on some phenotypes, adjusting for them does not impact PGI estimates in any meaningful way.

#### Table S3: P-values of comparisons of PGI coefficients before and after adding PCs to within-family models

| | Base vs. 4 PCs | 4 PCs vs. 8 PCs | 8 PCs vs. 12 PCs | 12 PCs vs. 16 PCs | 16 PCs vs. 20 PCs | 20 PCs vs. 24 PCs | 24 PCs vs. 28 PCs | 28 PCs vs. 32 PCs | 32 PCs vs. 36 PCs | 36 PCs vs. 40 PCs | Base vs. 40 PCs |
|---|---|---|---|---|---|---|---|---|---|---|---|
| BMI | 0.9825 | 0.9982 | 0.9937 | 0.9963 | 0.9927 | 0.9852 | 0.9944 | 0.9949 | 0.9922 | 0.9870 | 0.9579 |
| Height | 0.9414 | 0.9990 | 0.9889 | 0.9968 | 0.9957 | 0.9966 | 0.9979 | 0.9974 | 0.9931 | 0.9995 | 0.9558 |
| Educational Attainment | 0.9832 | 0.9932 | 0.9837 | 0.9974 | 0.9968 | 0.9828 | 0.9985 | 0.9886 | 0.9997 | 0.9997 | 0.9530 |
| Number of Children Ever Born | 0.9971 | 0.9875 | 0.9914 | 0.9943 | 0.9997 | 0.9959 | 0.9950 | 0.9925 | 0.9974 | 0.9959 | 0.9986 |
| Cigarettes Per Day | 0.9894 | 0.9865 | 0.9980 | 0.9967 | 0.9970 | 0.9992 | 0.9989 | 0.9984 | 0.9938 | 0.9990 | 0.9717 |
| Drinks Per Week | 0.9753 | 0.9977 | 0.9907 | 0.9991 | 0.9995 | 0.9928 | 0.9953 | 0.9997 | 0.9943 | 0.9978 | 0.9820 |
| Diastolic BP | 0.9844 | 0.9973 | 0.9922 | 0.9939 | 0.9979 | 0.9975 | 0.9947 | 0.9827 | 0.9996 | 0.9894 | 0.9582 |
| Seen Doc/GP for Depression | 0.9949 | 0.9987 | 0.9927 | 0.9955 | 0.9960 | 0.9914 | 0.9914 | 0.9914 | 0.9941 | 0.9969 | 0.9939 |
| Birthweight | 0.9893 | 0.9977 | 0.9974 | 0.9910 | 0.9934 | 0.9936 | 0.9995 | 0.9932 | 0.9997 | 0.9999 | 0.9996 |

P-values are reported from tests comparing PGI estimates (constructed using the largest available population GWAS estimates) before and after the inclusion of proband PCs. Models adjust for the relevant parental PGI, age, sex and array.We used the method for comparing regression coefficients reported in Clogg, Petkova, and Haritou (1995). None of the comparisons are statistically significant.

##### *Constructing PGI using GWAS conducted in unrelated sample of the UKB*

As a sensitivity check, we ran GWASs in the unrelated white British sample of the UKB for the nine phenotypes using the same procedure across all phenotypes. Sex, age, array, and 40 PCs calculated in the white British subsample were included as controls in the GWAS. Standard quality controls measures were performed prior to GWAS: we included SNPs with MAF of at least 5%, excluded individuals with more than 10% missingness and with discordant reported and biological sex, and excluded SNPs that violate HWE test with p-value of <1e-6.

We then ran models where we included mid-parent and proband PGI for a given phenotype, and we tested whether the coefficient changed once we included proband PCs in bundles of 4. Table

S4 reports the p-values of regression coefficient comparison tests (Clogg, Petkova, and Haritou 1995). None of the comparisons are statistically significant.

**Table S4: P-values from comparisons of PGI coefficients before and after adding PCs to within-family models (self-constructed GWAS)**

| | Base vs. 4 PCs | 4 PCs vs. 8 PCs | 8 PCs vs. 12 PCs | 12 PCs vs. 16 PCs | 16 PCs vs. 20 PCs | 20 PCs vs. 24 PCs | 24 PCs vs. 28 PCs | 28 PCs vs. 32 PCs | 32 PCs vs. 36 PCs | 36 PCs vs. 40 PCs | Base vs. 40 PCs |
|---|---|---|---|---|---|---|---|---|---|---|---|
| BMI | 0.999 | 0.988 | 0.999 | 0.995 | 1.000 | 0.995 | 0.982 | 0.998 | 0.994 | 0.996 | 0.960 |
| Height | 0.935 | 0.982 | 0.991 | 0.991 | 0.995 | 0.993 | 0.996 | 0.972 | 0.999 | 0.998 | 0.883 |
| Educational Attainment | 0.963 | 0.985 | 0.983 | 0.986 | 0.998 | 0.975 | 0.993 | 0.978 | 0.999 | 0.999 | 0.866 |
| Number of Children Ever Born | 0.997 | 0.975 | 0.995 | 0.999 | 0.983 | 0.996 | 0.997 | 0.992 | 0.998 | 0.990 | 0.973 |
| Cigarettes Per Day | 0.963 | 0.993 | 0.998 | 0.981 | 0.995 | 0.994 | 0.991 | 0.999 | 1.000 | 1.000 | 0.931 |
| Drinks Per Week | 0.993 | 0.979 | 0.999 | 0.988 | 0.993 | 0.998 | 0.998 | 0.985 | 0.991 | 1.000 | 0.926 |
| Diastolic BP | 0.966 | 0.995 | 0.984 | 0.995 | 0.997 | 0.997 | 0.983 | 1.000 | 0.998 | 0.993 | 0.980 |
| Seen Doc/GP for Depression | 0.978 | 0.995 | 0.991 | 0.998 | 0.998 | 0.999 | 0.981 | 0.999 | 0.989 | 0.990 | 0.994 |
| Birthweight | 0.990 | 0.993 | 0.996 | 0.991 | 0.998 | 0.998 | 0.998 | 1.000 | 0.995 | 0.999 | 0.985 |

P-values are reported from tests comparing PGI estimates (constructed using GWAS estimates from the white British unrelated sample) before and after the inclusion of proband PCs. Models adjust for the relevant parental PGI, age, sex and array. We used the method for comparing regression coefficients reported in Clogg, Petkova, and Haritou (1995). None of the comparisons are statistically significant.

**Figure S1: First two principal components and country of origin in the UK Biobank White British subset**

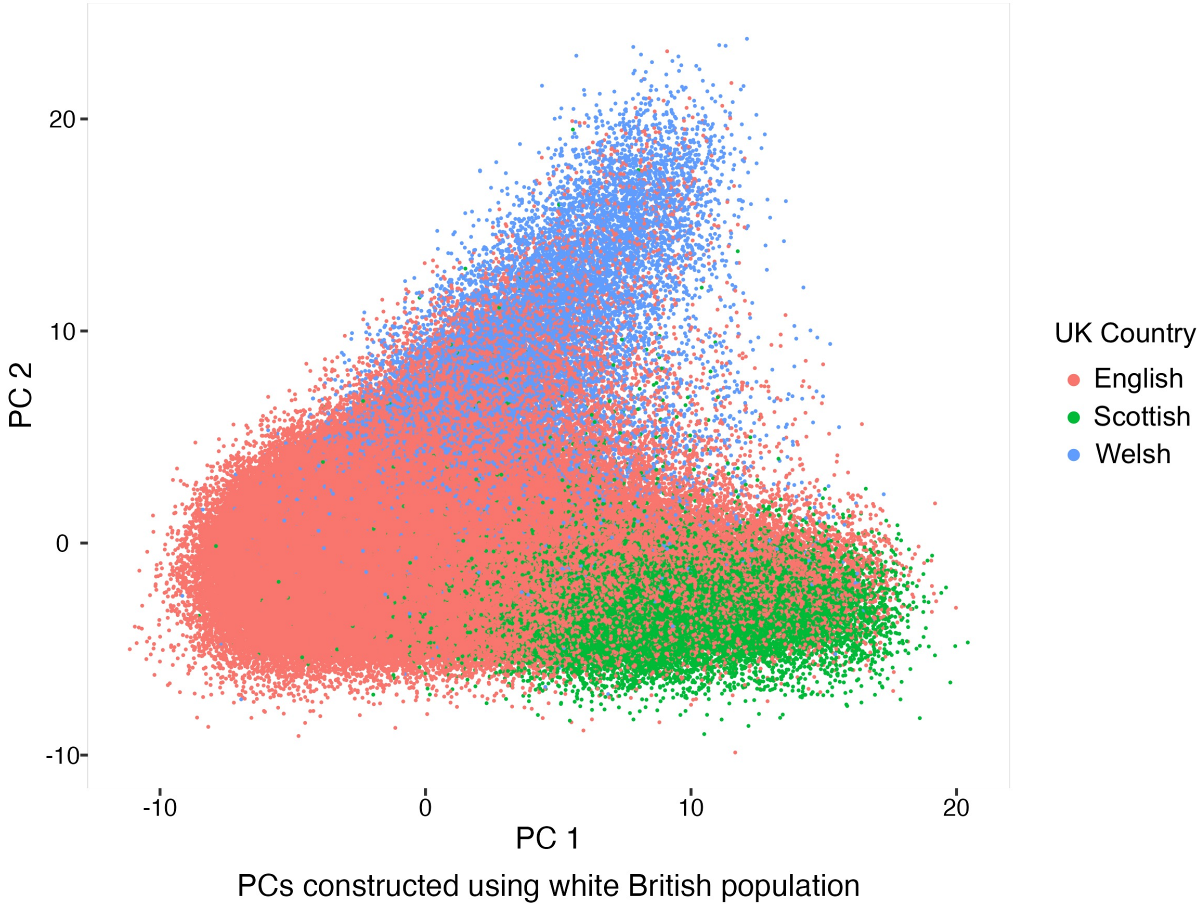


Scatter plot of PC1 and PC2 constructed using the white British population. The dots represent white British individuals. The colors represent different Great Britain Countries (England, Scotland, and Wales) based on respondents' place of birth. PC1 generally distinguishes between English and Scottish individuals, and PC2 Welsh and non-Welsh individuals.

### Figure S2: Comparing within-sibship effects of 40 PCs to their population effects by phenotype

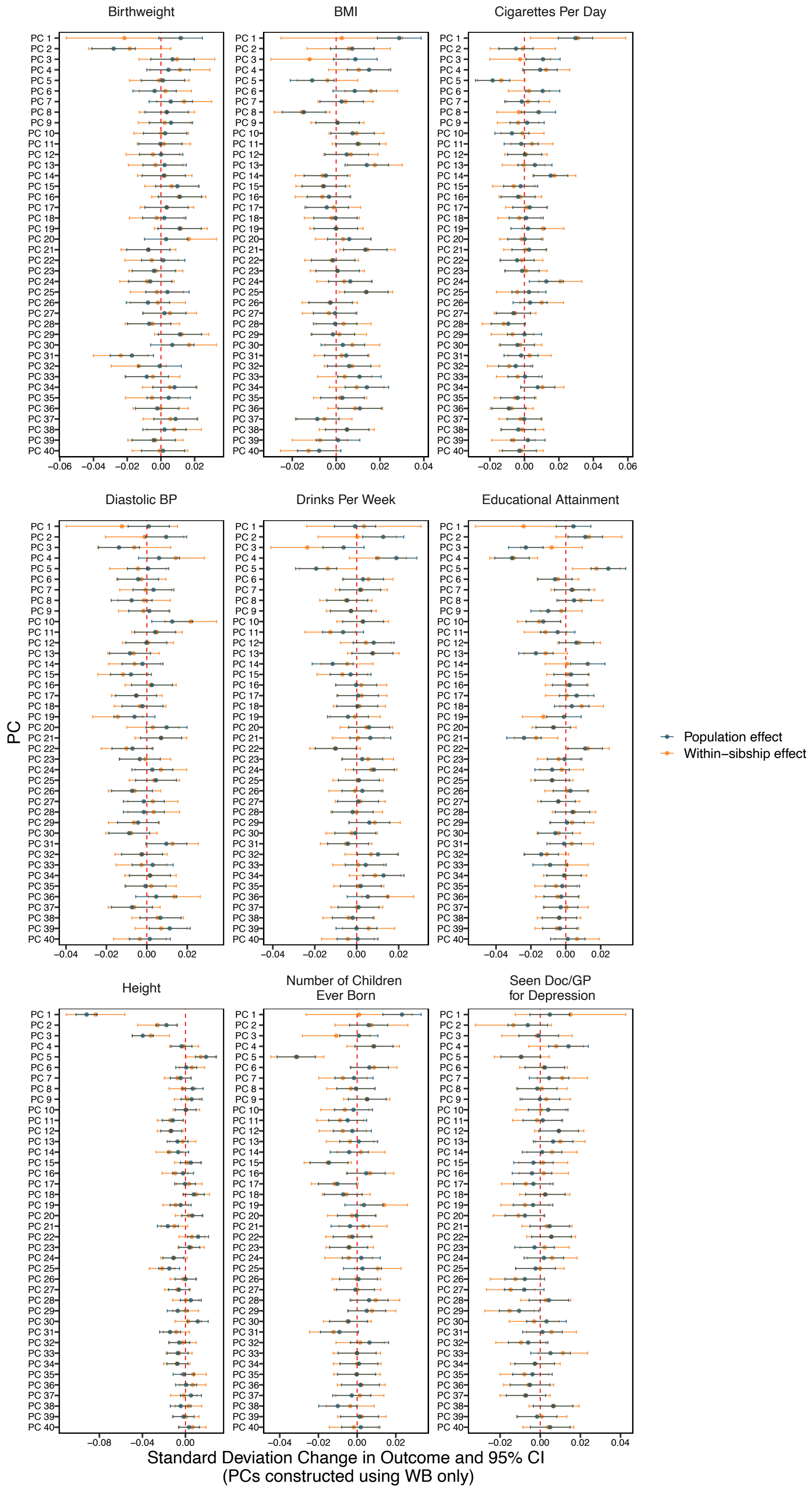

Standard deviation change in phenotype by PCs (point estimate ∓ 95% confidence interval) are plotted in population models (black) and in within-family models (orange). The nine panels correspond to the nine different phenotypes included in this study. The population models include controls and all 40 proband PCs and use the unrelated White British subsample. The within-family models include the sibling subsample and include controls, all 40 proband PCs, and all 40 parental PCs.
